# Gastrin-releasing peptide signaling in the nucleus accumbens medial shell regulates neuronal excitability and motivation

**DOI:** 10.1101/2024.05.29.596538

**Authors:** Erin E. Aisenberg, Thomas L. Li, Hongli Wang, Atehsa A. Sahagun, Emilie M. Tu, Helen S. Bateup

## Abstract

Neuropeptides are the largest class of neuromodulators. It has been shown that subpopulations of dopamine neurons express mRNA for the neuropeptide Gastrin-releasing peptide (GRP); however, its functional relevance in dopaminergic circuits is unknown. Here, we find that the GRP receptor (GRPR) is present in the nucleus accumbens medial shell (NAc MSh), which is targeted by GRP-expressing midbrain dopamine neurons as well as glutamatergic inputs from the hippocampus and amygdala. We show that the NAc MSh GRPR-positive cells are a subpopulation of D2 receptor-expressing neurons, comprising both classical indirect pathway striatal projection neurons (iSPNs) and eccentric SPNs (eSPNs), which have high intrinsic excitability, and can be activated by GRP *in vivo*. NAc-specific deletion of *Grpr* increases motivation in a progressive ratio test, demonstrating that GRPR regulates motivated behaviors. These experiments establish GRP/GRPR signaling as a potent modulator of mesolimbic circuits and advance our understanding of neuropeptide actions in the brain.

## Introduction

Neuromodulators play a critical role in the nervous system via their regulation of neuronal excitability, synaptic transmission, and plasticity^1, 2^. The largest family of known neuromodulators are neuropeptides—small short-chain polypeptides that act via metabotropic G-protein-coupled receptors (GPCRs)^3^. Compared to classical neurotransmitters, neuropeptides lead to diverse effects on their target neurons and regulate numerous physiological functions such as eating, sleeping, sexual behaviors, stress response, and pain perception^4, 5^. Additionally, neuropeptides have distinct machinery for their synthesis and transport and cannot be synthesized or recycled in axons^6^. It has recently become appreciated that many neurons co-express genes for both neurotransmitters and one or more neuropeptides^7, 8^. In particular, a recent single-cell RNA-sequencing (scRNA-seq) study found that 98% of mouse cortical neurons express at least one neuropeptide or neuropeptide receptor gene^7^. As such, neuropeptides are poised to make a major contribution to brain function and behavior; however, an understanding of their breadth, scope and mechanisms of action is currently limited.

One such neuropeptide is Gastrin-releasing peptide (GRP), a bombesin-like neuropeptide that signals through a Gq-coupled GPCR (gastrin-releasing peptide receptor, GRPR, also called BB2)^9,10^. GRP was originally named for its role in stimulating gastrin release from G cells in the stomach^11^. In the central nervous system, GRP has been studied in the spinal cord in the context of itch and pain processing^12–14^. In the brain, several studies have demonstrated a role for GRP/GRPR signaling in emotionally-motivated and fear memories as well as responses to stress^15, 16^. In the lateral nucleus of the amygdala, GRP signaling regulates fear^17^. In this region, GRP excites the GABAergic interneurons that express GRPR, increasing their inhibition of the principle neurons expressing *Grp* via a negative feedback mechanism^17^. In the auditory cortex, *Grpr* is expressed in vasoactive intestinal polypeptide (VIP) neurons and GRP signaling recruits disinhibitory cortical microcircuits and enhances auditory fear memories^18^. At the cellular level, GRP wash-on in brain slices leads to depolarization, and in some cases, rhythmic or tonic firing, in a variety of neuron types^4, 19–21^.

A recent scRNA-seq study of mouse midbrain dopamine (DA) neurons found that 30% of midbrain DA neurons in the ventral tegmental area (VTA) and the ventral substantia nigra pars compacta (SNc) express mRNA coding for *Grp*^8^, consistent with prior studies demonstrating GRP immunoreactivity in the SNc^22^. Additionally, *Grp* has been identified as a differentially expressed gene between VTA and SNc DA neurons^23, 24^. *GRP*-expressing DA neurons have also been observed in post-mortem human brains and are suggested to be relatively resistant to degeneration in Parkinson’s disease^23^. When examining the projection targets of the *Grp*+ DA neurons, Kramer et al. (2018) found two populations, one that resides in the ventral SNc and projects to the dorsal medial striatum and the other that is in the medial VTA and targets the nucleus accumbens medial shell (NAc MSh)^8^. These findings suggest that GRP may be co-released by DA neurons and may be a functionally relevant signaling molecule in the striatum.

The striatum is the main input structure of the basal ganglia, which integrates glutamatergic inputs from the cortex and thalamus, DA from the midbrain, GABAergic inhibition from multiple intra-striatal sources, and acetylcholine from cholinergic interneurons^25^. GABAergic striatal projection neurons (SPNs) are the major output neuron of the striatum, comprising 90-95% of the neuronal population^26^. SPNs are canonically divided into direct-pathway and indirect-pathway neurons based on their primary projections to either the substantia nigra pars reticulata and globus pallidus internal segment (SNr/GPi) or globus pallidus external segment (GPe), respectively^27^. Direct pathway SPNs (dSPNs) typically express type 1 DA receptors (D1) while indirect pathway SPNs (iSPNs) generally express type 2 DA receptors (D2)^27^. A third type of SPN, eccentric SPNs (eSPNs) was recently proposed^28^. These eSPNs are defined by their expression of SPN markers including *Ppp1r1b*, *Adora2a*, and *Drd1* as well as a unique gene module consisting of ∼100 genes^28^. Unlike dSPNs and iSPNs, little is known abuot the functional properties of eSPNs.

The dorsal striatum (DS) is important for motor functions including action selection and motor learning, while the ventral striatum controls motivated behaviors and reward learning^29^. In the DS, activation of dSPNs with optogenetics promotes locomotor activity while bulk activation of iSPNs inhibits movement^30, 31^. During normal movement, however, both pathways are active and must work in a coordinated manner^32, 33^. The ventral striatum (nucleus accumbens, NAc) also contains direct- and indirect-pathway SPNs. In the NAc, the dSPNs have canonically been thought to directly project to and inhibit the VTA, promoting motivated behavior through disinhibition of VTA GABA neurons that synapse onto VTA DA neurons and project back to the NAc^34^. NAc iSPNs first project to, and inhibit the ventral pallidum (VP); VP neurons then project to the VTA and ultimately the thalamus, resulting in inhibition of motivated behaviors^34^.

With increasing use of cell type-specific approaches, it has become clear that the canonical divisions of the striatum: D1 vs D2 SPNs, dorsal vs ventral striatum, indirect vs direct pathway are oversimplified and there is likely further complexity to uncover within these circuits^35, 36^. While it is well known that neuropeptides including opioids are active within the striatum, opioid receptors are Gi/o-coupled, and the opioids themselves are synthesized within the striatum^3^. In contrast, the *Grp*+ DA neurons described above would bring in a neuropeptide from outside the striatum and are hypothesized to increase the excitability of GRPR-expressing neurons.

The goal of this study was to assess a possible role for GRP-GRPR signaling in the NAc. We find that *Grpr* is expressed in the NAc medial shell (MSh), and *Grpr*-expressing cells have unique electrophysiological properties. Specifically, a majority of *Grpr*+ cells are spontaneously active in slice recordings, and have high intrinsic excitability. Through fluorescent *in situ* hybridization (FISH) and scRNA-seq data from the Allen Institute, we identify *Grpr*+ cells as mainly D2R-expressing SPNs^37^. Further, we find multiple distinct sub-populations of *Grpr*+ cells, both canonical iSPNs as well as eSPNs, based on gene expression patterns and electrophysiological signatures. In addition to the *Grp*+ DA neurons, we show that the NAc MSh receives glutamatergic *Grp*+ inputs from the hippocampus and amygdala. GRP activates and increases the excitability of GRPR neurons in the NAc MSh both in slice and *in vivo*. Finally, we demonstrate that specific deletion of *Grpr* in the NAc MSh increases motivation in a progressive ratio test. These findings identify previously uncharacterized cell types in the NAc MSh, provide functional characterization of eSPNs and comparison to classic SPNs, and map out a GRP-GRPR circuit that mediates striatal excitability and motivated behaviors.

## Results

### Grpr is expressed in striatal neurons

To investigate whether GRP may be a functionally relevant neuromodulator in nigrostriatal and mesolimbic DA circuits, we used fluorescent *in situ* hybridization (FISH) to probe for expression of mRNA encoding *Grpr* in the DS and NAc MSh (Fig. 1a). Indeed, we found *Grpr* expression in subpopulations of cells in both the DS and NAc MSh (Fig. 1b). These cells were concentrated in the anterior portion of the striatum, with the highest concentration of *Grpr*+ cells found around A/P +1.18 mm from bregma (Fig. 1b). Our findings are consistent with prior autoradiographic studies showing a high density of binding sites for bomesin-like peptides in the NAc and striatum^38, 39^. Given the fact that the NAc was previously identified as a region with high GRPR expression^38, 39^, we focused further analysis on the *Grpr*-expressing cell population in the NAc MSh, which receives dopaminergic inputs from the VTA.

**Fig. 1:**
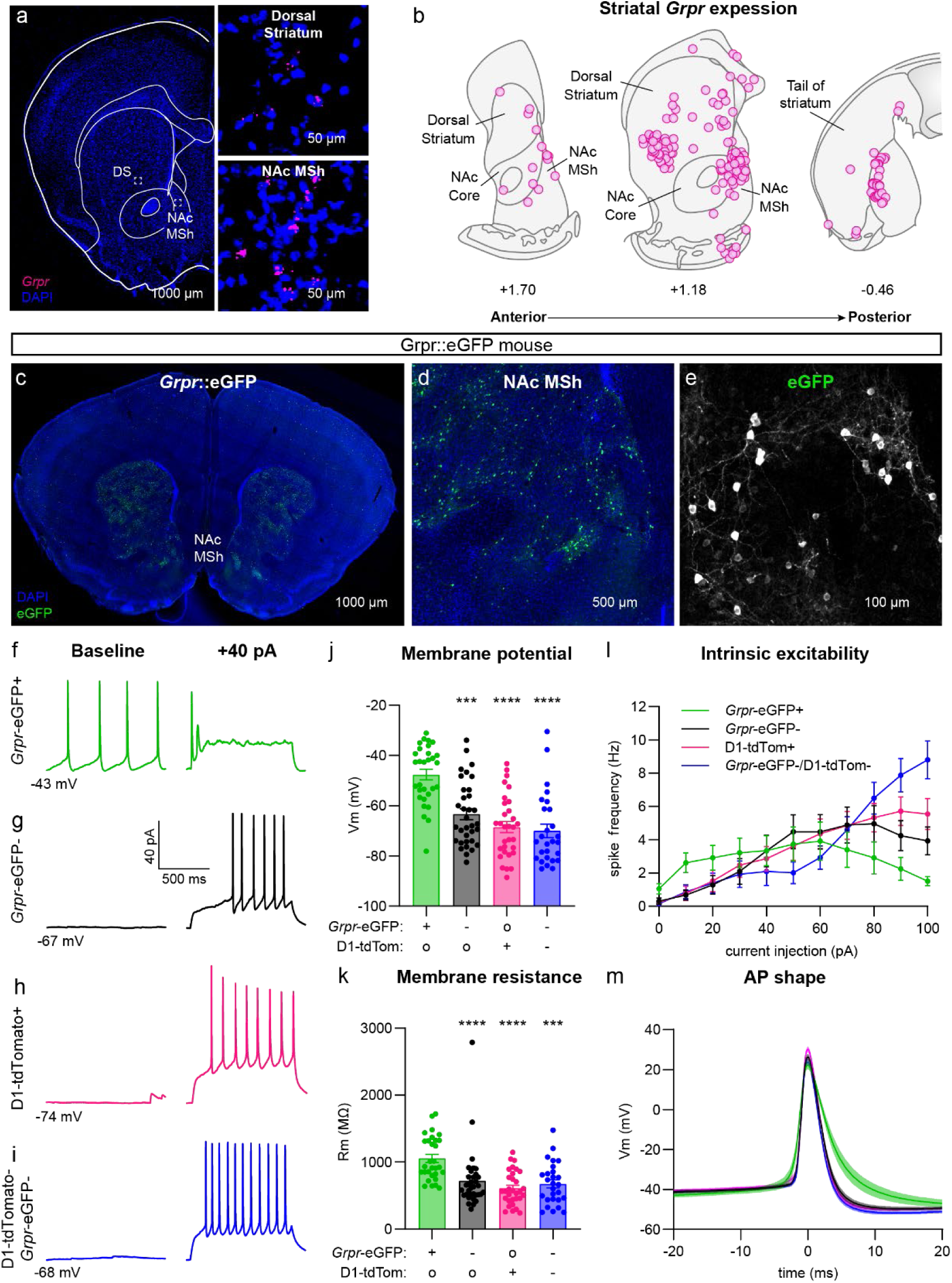
*Grpr* is expressed in the dorsal and ventral striatum and *Grpr*+ cells in the NAc MSh and have unique physiological properties. **(a)** Representative image of a coronal brain section (A/P +1.18) from a wild-type (WT) mouse. Fluorescent *in situ* hybridization for *Grpr* in magenta. Nuclei are labeled in blue with DAPI. Insets show higher magnification images for dorsal striatum (DS) and nucleus accumbens medial shell (NAc MSh). **(b)** Summary schematic of *Grpr* expression across the A/P axis of the DS and NAc MSh. Each dot represents the location of a *Grpr*+ cell. Dots are summed across 3 WT mice. **(c)** Representative image of a coronal brain section (A/P +1.18) from a *Grpr*::eGFP mouse. eGFP in green, DAPI labeled nuclei in blue. **(d)** Zoom-in of the NAc MSh from panel **c**. **(e)** Zoom-in of panel **d** showing eGFP+ cells in grayscale. **(f**-**i)** Representative traces from (f) *Grpr-*eGFP+, (g) *Grpr-*eGFP-, **(h)** D1-tdTomato+, and **(i)** D1-tdTomato-/*Grpr*-eGFP- (presumed D2+) cells in the NAc MSh recorded at baseline (left) and following +40 pA current injection (right). **(j)** Mean ± SEM membrane potential of NAc MSh neurons. *Grpr*-eGFP+ in green: n=30 cells from 15 mice, *Grpr*- eGFP- in black: n=34 cells from 11 mice, D1-tdTomato+ in magenta: n=31 cells from 5 mice, and D1-tdTomato-/*Grpr*-eGFP-in dark blue: n=27 cells from 3 mice. (****p<0.0001, one-way ANOVA; Kruskal-Wallis with Dunn’s correction for multiple comparisons, ***p=0.0003 *Grpr*-eGFP+ vs *Grpr*- eGFP-, ****p<0.0001 *Grpr*-eGFP+ vs D1-tdTomato+, ****p<0.0001 *Grpr*-eGFP+ vs D1-tdTomato-/*Grpr*-eGFP-). **(k)** Mean ± SEM membrane resistance of NAc MSh neurons. n is the same as panel **j**. (****p<0.0001, one-way ANOVA; Kruskal-Wallis with Dunn’s correction for multiple comparisons, ****p<0.0001 *Grpr*-eGFP+ vs *Grpr*-eGFP-, ****p<0.0001 *Grpr*-eGFP+ vs D1-tdTomato+, ***p=0.0001 *Grpr*-eGFP+ vs D1-tdTomato-/*Grpr*-eGFP-). **(l)** Input-output curves showing the mean ± SEM firing frequency of NAc MSh neurons in response to positive current steps of increasing amplitude. *Grpr*-eGFP+ in green: n=23 cells from 10 mice, *Grpr*-eGFP- in black: n=21 cells from 7 mice, D1-tdTomato+ in magenta: n=31 cells from 5 mice, and D1-tdTomato-/*Grpr*-eGFP- in dark blue: n=27 cells from 3 mice. (Two-way repeated measures ANOVA, current injection x cell type ****p<0.0001, current injection ****p<0.0001, cell type p=0.7206) **(m)** Mean ± SEM membrane potential over time for the first action potential (AP) evoked by current injection in each cell. *Grpr*-eGFP+ in green: n=18 cells from 10 mice, *Grpr*- eGFP- in black: n=20 cells from 7 mice, D1-tdTomato+ in magenta: n=27 cells from 5 mice, and D1-tdTomato-/*Grpr*-eGFP- in dark blue: n=25 cells from 3 mice. For panels **j** and **k**, + indicates the presence of a fluorophore labeling the given cell type, - indicates that the cell was negative for the fluorophore, and o indicates that the tissue did not have a fluorophore that would label the given cell type.

To study the properties of GRPR-expressing cells in the NAc MSh, we obtained a transgenic mouse line that fluorescently labels GRPR+ cells with eGFP (Fig. 1c-e). *Grpr*::eGFP mice (Gensat [MMRRC #036178-UCD])^40^ have previously been shown to express eGFP exclusively in *Grpr*+ cells^12^. To characterize the morphology of GRPR+ cells in the NAc MSh, we filled cells with neurobiotin through a patch pipette and performed cellular reconstructions of eGFP+ and eGFP-cells (Extended Data Fig. 1a,b). We found that GRPR+ cells had similar dendritic complexity as neighboring cells, the majority of which are expected to be SPNs (Extended Data Fig. 1c). Additionally, we observed no differences in the total dendritic arbor length or the number of primary dendrites in eGFP+ vs eGFP-cells (Extended Data Fig. 1d,e). We also examined dendritic spines in cells transduced with an mCherry-expressing virus and observed that GRPR+ cells were spiny and had a similar spine density as neighboring neurons (Extended Data Fig. 1f-h). This analysis suggests that GRPR+ cells in the NAc MSh are likely a subpopulation of SPNs, as striatal interneurons have few or no spines^41^.

We then bred the *Grpr*::eGFP mice to Drd1-tdTomato mice^42^ to determine whether the GRPR+ cells belonged to a given SPN subtype (Extended Data Fig. 1i-k). We noted that in the NAc MSh, the eGFP+ cells had little overlap with the tdTomato+ cells (∼5%) (Extended Data Fig. 1k). This, along with the presence of dendritic spines, suggested that the GRPR+ cells in the NAc MSh were likely a subpopulation of D2-expressing SPNs.

We next assessed the functional and physiological properties of GRPR-expressing neurons using the *Grpr*::eGFP;D1-tdTomato mice^42^ which allowed us to visualize both GRPR+ cells and dSPNs and we could deduce the unlabeled cells were likely GRPR-iSPNs given the cell type proportions in the NAc MSh. We used whole-cell electrophysiology to record from *Grpr-*eGFP+ cells and compared them to neighboring *Grpr-*eGFP-neurons in the NAc MSh (Fig. 1f,g). We also recorded from genetically identified dSPNs (D1-tdTomato+, Fig. 1h) and presumed iSPNs that were GRPR negative (D1-tdTomato-;*Grpr*-eGFP-, Fig. 1i). Interestingly, we found that the baseline physiological properties of *Grpr*-eGFP+ cells differed significantly from other SPNs in the NAc MSh. Approximately 75% of the *Grpr*-eGFP+ cells were spontaneously active at baseline (Fig. 1f), which was not characteristic of eGFP-cells or known SPN subtypes (Fig. 1g-i). We also found that compared to neighboring cell populations, *Grpr*-eGFP+ neurons were significantly more depolarized (Fig. 1j) and had higher membrane resistance (Fig. 1k). *Grpr*-eGFP+ cells also differed in their input-output curve compared to neighboring SPNs, exhibiting increased action potential firing in response to small positive currents and a transition to depolarization block with larger currents (Fig. 1l). We also noted the *Grpr*-eGFP+ cells differed in the shape of their action potentials, particularly in terms of width (*Grpr*-eGFP+ 5.2970 ± 0.4821 ms vs. Grpr-eGFP-3.2027 ± 0.1830 ms vs. D1-tdTomato+ 3.0003 ± 0.1076 ms vs. *Grpr*-eGFP-/D1-tdTomato-2.9597 ± 0.1365 ms; ****p<0.0001, one-way ANOVA; Holm-Sidak’s multiple comparison tests, ****p<0.0001 *Grpr*-eGFP+ vs *Grpr*-eGFP-, ****p<0.0001 *Grpr*-eGFP+ vs D1-tdTomato+, ****p<0.0001 *Grpr*-eGFP+ vs D1-tdTomato-/*Grpr*-eGFP-, Fig. 1m).

To determine whether expression of GRPR contributed to the high baseline excitability of *Grpr*-eGFP+ cells, we compared the membrane potential and resistance of *Grpr*-eGFP+ cells with and without DPDMB, a GRPR-specific antagonist. We found a small decrease in the membrane resistance of *Grpr*-eGFP+ cells treated with DPDMB compared to untreated cells (Extended Data Fig. 2a-c). This suggests that GRPR either has basal activity that can be blocked with DPDMB or that there may be tonic GRP-GRPR signaling in the slice preparation that increases the excitability of *Grpr*-eGFP+ cells.

Given that *Grpr*-eGFP+ cells were spontaneously active and had properties that were distinct from other SPNs in the area, we assessed how they compared to cholinergic interneurons, which are known to be tonically active^43^ (Extended Data Fig. 3). We recorded from tdTomato-labeled neurons in *Chat*-Cre;Ai9 mice^44, 45^ and found that while the *Chat*-Cre+ cells tended to be spontaneously active (53% of cells were firing at baseline, Extended Data Fig. 3b), their membrane potential and membrane resistance were significantly different from *Grpr*-eGFP+ cells (Extended Data Fig. 3c,d). Additionally, with increasing positive current injections, the *Chat+* cells were able to continue to fire action potentials, while the *Grpr*-eGFP+ cells went into depolarization block with >60 pA current injection (Extended Data Fig. 3e). This, together with the fact that ChAT+ cells are known to be aspiny^46–48^, indicated that GRPR+ cells are likely not a type of cholinergic interneuron.

### Grpr-expressing neurons in the NAc MSh are SPNs

Since *Grpr*-expressing cells had non-canonical properties compared to other neurons in the NAc MSh, we used multi-plex FISH with markers for known striatal cell types to determine their identify (Fig. 2a-j). We found that the vast majority of *Grpr*+ cells in the NAc MSh were positive for *Gad2* mRNA (96.3%) (Fig. 2a,b), and no *Grpr*+ cells were *Chat*+ (Fig. 2c,f), indicating that *Grpr*+ cells are GABA-ergic and not cholinergic. In the DS, we found similar expression patterns with 92.7% of *Grpr*+ cells expressing *Gad2*+ (Extended Data Fig. 4a,b) compared to only 1.9% co-expressing *Chat* mRNA (Extended Data Fig. 4c,f). To determine whether the *Grpr*+ cells were SPNs or a type of GABAergic interneuron, we probed for *Ppp1r1b*, which encodes the SPN-marker DARPP-32 (Fig. 2c,g). Nearly all *Grpr*-expressing cells co-expressed *Ppp1r1b* in the NAc MSh (93.8%) (Fig. 2g) and DS (75.5%) (Extended Data Fig. 4c,g), indicating that they were likely SPNs.

**Fig. 2:**
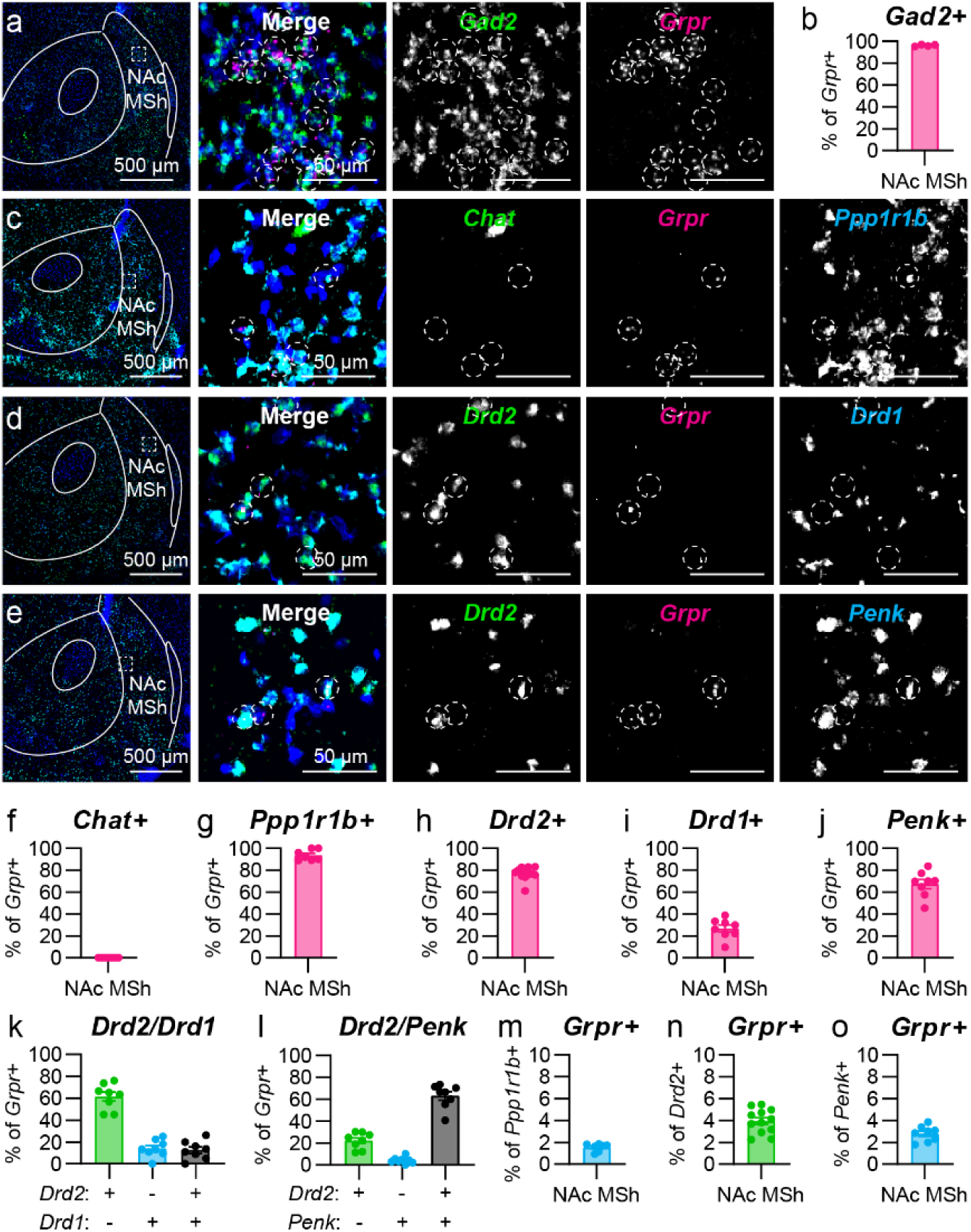
*Grpr* is expressed in SPNs in the NAc MSh. **(a)** Representative image of the NAc MSh. Fluorescent *in situ* hybridization (FISH) for *Gad2* in green and *Grpr* in magenta. Nuclei labeled with DAPI in blue. Dashed circles outline *Grpr+* cells. **(b)** Mean ± SEM percentage of *Grpr*+ cells in the NAc MSh and that are *Gad2+* in WT mice. (n=4 mice). **(c-e)** Representative images of NAc MSh. Nuclei labeled with DAPI in blue. Dashed circles outline *Grpr+* cells. **(c)** FISH for *Chat* in green, *Grpr* in magenta, and *Ppp1r1b* in cyan. **(d)** FISH for *Drd2* in green, *Grpr* in magenta, and *Drd1* in cyan. **(e)** FISH for *Drd2* in green, *Grpr* in magenta, and *Penk* in cyan. **(f-j)** Mean ± SEM percentage of *Grpr*+ cells in the NAc MSh of WT mice that are **(f)** *Chat+* (n=7 mice), **(g)** *Ppp1r1b*+ (n=7 mice), **(h)** *Drd2*+ (n=12 mice), **(i)** *Drd1*+ (n=8 mice), or **(j)** *Penk*+ (n=8 mice). **(k)** Mean ± SEM percentage of *Grpr*+ cells in the NAc MSh that are *Drd2*+/*Drd1*-(green), *Drd2*-/*Drd1*+ (blue), or *Drd2*+/*Drd1*+ (black) in WT mice (n=8 mice). **(l)** Mean ± SEM percentage of *Grpr*+ cells in the NAc MSh that are *Drd2*+/*Penk*- (green), *Drd2*-/*Penk*+ (blue), or *Drd2*+/*Penk*+ (black) in WT mice (n=8 mice). **(m-o)** Mean ± SEM percentage of **(m)** *Ppp1r1b*+ (n=7 mice), **(n)** *Drd2*+ (n=12 mice), or **(o)** *Penk*+ cells (n=8 mice) in the NAc MSh that are *Grpr*+.

To determine the specific SPN identity of the *Grpr*+ cells, we probed for mRNA encoding D1 (*Drd1*) and D2 (*Drd2*) DA receptors (Fig. 2d), which are typically expressed in dSPNs and iSPNs, respectively. The majority of *Grpr*+ cells in the NAc MSh expressed *Drd2* mRNA (77.3%) (Fig. 2h), suggesting that they were iSPNs. However, a small subpopulation of *Grpr*+ cells in the NAc MSh expressed *Drd1* (26.7%) (Fig. 2i). Multiple studies have reported that a proportion (5-30%) of SPNs in the NAc MSh express both *Drd1* and *Drd2*^35, 49, 50^; therefore, we examined whether the *Grpr*+/*Drd1+* population in the NAc MSh were dSPNs or SPNs co-expressing both DA receptors. We found that 61.5% of *Grpr*+ cells in the NAc MSh expressed *Drd2*, but not *Drd1*, 14.1% expressed *Drd1* only, and 12.6% expressed both *Drd1* and *Drd2* (Fig. 2k). Given the preferential expression of *Drd2* in *Grpr*+ cells, we probed for the iSPN marker *Penk* (Fig. 2e) and found that it was expressed by the majority (67.4%) of *Grpr*+ cells in the NAc MSh (Fig. 2j). We also examined the co-localization between *Drd2* and *Penk* mRNA and found that 22.2% of the *Grpr*+ cells in the NAc MSh expressed *Drd2*, but not *Penk*, 4.3% expressed *Penk*, but not *Drd2*, and 63.1% of *Grpr*+ cells in the NAc MSh expressed both markers (Fig. 2l). Together, this analysis indicates that *Grpr*+ cells in the NAc MSh are likely to be GABA-ergic D2R-expressing SPNs. We note that only a small percentage of all *Prpp1r1b* (1.5%), *Drd2* (3.9%), or *Penk*-expressing (2.8%) cells in the MSh were *Grpr* positive, indicating that *Grpr*+ cells are a select subpopulation of SPNs in the NAc MSh (Fig. 2m-o).

We next examined the SPN identity of *Grpr*+ cells in the DS (Extended Data Fig. 4). While a majority of *Grpr*+ cells in the DS expressed SPN markers (Extended Data Fig. 4c-j), the SPN sub-type classification was less clear. *Drd2* mRNA was expressed in 32.7% of DS *Grpr*+ cells, while *Drd1* mRNA was expressed in 43.3% of *Grpr*+ cells, (Extended Data Fig. 4h,i). We also found that *Penk* mRNA was found in 38.0% of DS *Grpr*+ cells, consistent with the proportion expressing *Drd2* mRNA (Extended Data Fig. 4j). Together this suggests that *Grpr* is expressed in both types of SPNs in the DS.

We were intrigued by the portion of *Grpr*+ cells that expressed mRNA for *Drd2*, but not the other canonical iSPN marker, *Penk*. We therefore made use of the whole mouse brain scRNA-seq atlas from the Allen Institute for Brain Science^37^ to further explore the molecular identity of *Grpr+* cells (Extended Data Fig. 5). We first extracted and clustered all of the cells in the mouse dorsal and ventral striatum (VS) (Extended Data Fig. 5a). We noted a clear separation between the DS and VS within the largest SPN clusters, defined by *Ppp1r1b* expression (Extended Data Fig. 5b,c). Feature plots highlighting *Ppp1r1b*, *Drd1*, *Drd2*, and *Grpr*-expressing cells were consistent with FISH results showing *Grpr*+ cells were mainly *Ppp1r1b*+/*Drd2*+ in the VS (Extended Data Fig. c-f).

We then subsetted the *Grpr*+ cells in the VS and clustered them into five distinct clusters (Extended Data Fig. 5g). The two largest clusters of *Grpr*-expressing cells (clusters 0 and 1) expressed *Gad2*, *Ppp1r1b*, and *Drd2*, but not *Drd1, Chat, Pvalb,* or *Sst* (Extended Data Fig. 5h-p), consistent with an iSPN identify. We noted that cluster 1 cells expressed *Drd2* but had lower expression of *Penk* (Extended Data Fig. 5l,m), reminiscent of the *Grpr+/Drd2+/Penk-* cells observed by FISH. We further explored the identify of cluster 1 cells and found that they expressed markers of eSPNs^28^ including *Casz1, Th, Otof, Cacng5,* and *Pcdh8* (Extended Data Fig. 5q-u). Together, this suggests that the majority of *Grpr+* cells in the VS are GABA-ergic *Drd2*-expressing SPNs that can be further divided into iSPN and eSPN sub-types based on their gene expression.

To test whether *GRPR*-expressing cells are also present in the human caudate/putamen, we applied a similar analysis to a scRNA-seq dataset from the human adult brain^51^ (Extended Fig. 6). Feature plots of *PPP1R1B*, *DRD1*, *DRD2*, and *GRPR* expression showed that in the human VS, *GRPR* is expressed in both *DRD1*- and *DRD2*-expressing SPNs (Extended Data Fig. 6b-e).

We isolated the *GRPR*-expressing cells in the VS and clustered them into four clusters (Extended Data Fig. 6f). In the human brain dataset, clusters 0, 1, and 2 expressed SPN markers, with cluster 0 corresponding to *DRD1*-expressing dSPNs, cluster 1 to *DRD2*-expressing iSPNs, and cluster 2 to *CASZ1-*expressing eSPNs (Extended Data Fig. 6f-t). Interestingly, unlike in the mouse dataset, there was also a population of *GRPR*+ *SST*-expressing cells, which may represent somatostatin interneurons (cluster 3, Extended Data Fig. 6o). Overall, while there was more diversity in the types of cells that expressed *GRPR* in the human VS, it was clear that *GRPR* is expressed in subpopulations of iSPNs and eSPNs, consistent with our observations in mice.

### Grpr+ cells in the NAc MSh comprise distinct physiological sub-types

Given that the scRNA-seq data had identified a population of *Grpr*+ cells in the VS that appeared to be eSPNs, we completed additional FISH experiments to probe for the eSPN marker *Casz1* in the NAc MSh (Fig. 3a-c). We found that 36.4% of *Grpr*+ cells in the NAc MSh expressed *Casz1* mRNA (Fig. 3b). This was notable given that 63.1% of *Grpr*+ cells in the NAc MSh expressed both *Drd2* and *Penk* (Fig. 2l); therefore, *Grpr+/Casz1*+ cells could represent the remaining fraction. This provides further evidence that there are two populations of *Grpr*+ cells in the NAc MSh, classical iSPNs and eSPNs. We also noted that not all eSPNs are *Grpr*+, as only 15.6% of *Casz1*+ cells in the NAc MSh were *Grpr*+ (Fig. 3c).

**Fig. 3:**
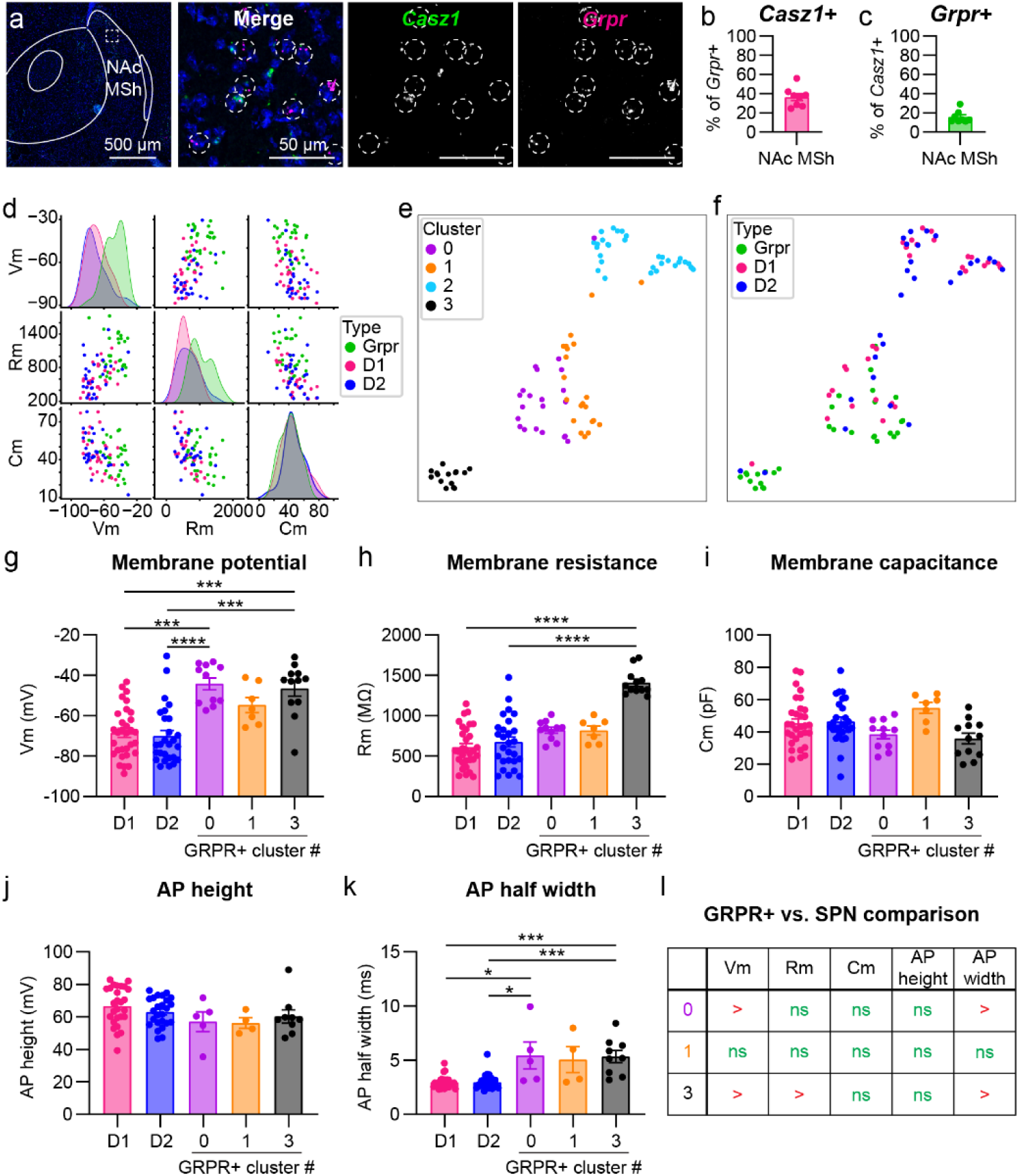
*Grpr*+ neurons in the NAc MSh can be clustered based on their physiological properties. **(a)** Representative image of the NAc MSh. FISH for *Casz1* in green and *Grpr* in magenta. Nuclei labeled with DAPI in blue. Dashed circles outline *Grpr+* cells. **(b)** Mean ± SEM percentage of *Grpr*+ cells in the NAc MSh and that are *Casz1+* in WT mice. (n=8 mice). **(c)** Mean ± SEM percentage of *Casz1*+ (n=8 mice) in the NAc MSh that are *Grpr*+. **(d)** Pair plot of passive properties from electrophysiology recordings from D1 (D1-tdTomato+), D2 (D1-tdTomato-/*Grpr*- eGFP-), and GRPR (*Grpr*-eGFP+) cells in NAc MSh. **(e)** UMAP of passive properties of D1, D2, and GRPR cells in NAc MSh. Vm, Rm, and Cm values for each cell were centered and scaled to unit variance, and UMAP embeddings were computed using n_neighbors=10 and min_dist=0. Clusters were determined using the K-Means algorithm with n_clusters=4. **(f)** UMAP of passive properties from **e** colored by cell type. **(g)** Mean ± SEM membrane potential of NAc MSh neurons. D1-tdTomato+ (D1) in magenta: n=31 cells from 5 mice, D1-tdTomato-/*Grpr*-eGFP-(D2) in dark blue: n=27 cells from 3 mice, *Grpr*-eGFP+ cells in cluster 0 in purple: n=11 cells from 7 mice, *Grpr*-eGFP+ cells in cluster 1 in orange: n=7 cells from 5 mice, and *Grpr*-eGFP+ cells in cluster 3 in black: n=12 cells from n=8 mice. (****p<0.0001, one-way ANOVA; Kruskal-Wallis with Dunn’s correction for multiple comparisons, ***p=0.0002 D1 vs 0, ****p<0.0001 D2 vs 0, p=0.2044 D1 vs 1, p=0.0951 D2 vs 1, ***p=0.0009 D1 vs 3, and ***p=0.0002 D2 vs 3). **(h)** Mean ± SEM membrane resistance of NAc MSh neurons. n is the same as for panel **g.** (****p<0.0001, one-way ANOVA; Kruskal-Wallis with Dunn’s correction for multiple comparisons, p=0.1880 D1 vs 0, p=0.6294 D2 vs 0, p=0.4887 D1 vs 1, p>0.9999 D2 vs 1, ****p<0.0001 D1 vs 3, and ****p<0.0001 D2 vs 3). **(i)** Mean ± SEM membrane capacitance of NAc MSh neurons. n is the same as for panel **g**. (*p=0.0169, one-way ANOVA; Kruskal-Wallis with Dunn’s correction for multiple comparisons, p>0.9999 D1 vs 0, p=0.5381 D2 vs 0, p=0.3473 D1 vs 1, p=0.7138 D2 vs 1, p=0.3684 D1 vs 3, and p=0.1607 D2 vs 3). **(j)** Mean ± SEM action potential (AP) height of NAc MSh neurons. D1-tdTomato+ (D1) in magenta: n=27 cells from 5 mice, D1-tdTomato-/*Grpr*-eGFP- (D2) in dark blue: n=25 cells from 3 mice, *Grpr*-eGFP+ cells in cluster 0 (0) in purple: n=5 cells from 4 mice, *Grpr*- eGFP+ cells in cluster 1 (1) in orange: n=4 cells from 3 mice, and *Grpr*-eGFP+ cells in cluster 3 (3) in black: n=9 cells from n=5 mice. (p=0.1230, one-way ANOVA; Kruskal-Wallis with Dunn’s correction for multiple comparisons, p=0.8731 D1 vs 0, p>0.9999 D2 vs 0, p=0.2781 D1 vs 1, p>0.9999 D2 vs 1, p=0.3553 D1 vs 3, and p>0.9999 D2 vs 3). **(k)** Mean ± SEM AP half-width of NAc MSh neurons. n is the same as for panel **j**. (****p<0.0001, one-way ANOVA; Kruskal-Wallis with Dunn’s correction for multiple comparisons, *p=0.0420 D1 vs 0, *p=0.0346 D2 vs 0, p=0.1229 D1 vs 1, p=0.1042 D2 vs 1, ***p=0.0005 D1 vs 3, and ***p=0.0004 D2 vs 3). **(l)** Summary table of **g-k**. Data for D1+ and D2+ cells in **g**-**i** is the same as in Figure 1, plotted here for comparison. Data for GRPR+ cells in **g**-**i** is the same as in Figure 1, but subdivided by cluster.

Given this new insight, we generated pair plots to examine how passive properties were distributed among GRPR+ SPNs and defined dSPNs (D1), and iSPNs (D2) (see Fig. 1). We found that for both membrane potential and resistance, the distribution for GRPR+ cells was shifted to higher values compared to other SPNs and there were two distinct peaks for the GPRR+ population (Fig. 3d). The distributions for capacitance were overlapping among all cell types, consistent with the similar morphological profile of GRPR+ and GRPR-SPNs in the NAc (see Extended Data Fig. 1). Because the pair plots indicated that there may be multiple distinct populations of GRPR+ cells, we performed dimensionality reduction and clustering analysis on our physiology data from Figure 1 (Fig. 3e,f). We used information about each cell’s membrane potential, resistance, and capacitance to generate a 2D UMAP embedding of the cells. Informed by this UMAP, we clustered the cells into 4 distinct clusters (Fig. 3e). GRPR+ cells belonged to three of these clusters: 0, 1, and 3 (Fig. 3f), indicating multiple distinct populations.

To delve into this further, we separated the GRPR+ cells based on their cluster identity and compared them to known D1+ and presumed D2+ cells in the NAc MSh (Fig. 3g-l). We examined membrane potential, resistance, and capacitance and action potential (AP) half-width and height (Fig. 3g-k). We found no significant differences in any of the examined properties between the SPN groups and the GRPR+ cells in cluster 1 (Fig. 3l). However, GRPR+ cells in clusters 0 and 3 had more depolarized membrane potential and greater AP half-width compared to canonical SPNs (Fig. 3h,k). Notably, cluster 0 and 3 cells differed in terms of their membrane resistance (Fig. 3h), with GRPR+ cells in cluster 3 having significantly higher membrane resistance than the SPN groups (Fig. 3h).

In both the *Casz1* FISH experiment and in the *Grpr*::eGFP mice, we noted that there were instances of high density “clumps” of *Grpr*+/eGFP+ cells in the ventral portions of the NAc MSh (Extended Data Fig. 7a-c). In the FISH experiments, *Grpr* and *Casz1* mRNA were highly co-expressed in these regions (Extended Data Fig. 7a). These clumps were reminiscent of immature cells migrating towards their final adult location^52^. We therefore hypothesized that some of the GRPR+ cells could be immature SPNs. To test this, we recorded from iSPNs in the NAc MSh from juvenile P8-P12 D2-GFP mice (Extended Data Fig. 7d-i). Cluster 0 and 1 GRPR+ cells had lower membrane resistance and higher capacitance compared to the immature cells, consistent with changes that occur in SPNs during postnatal maturation^53, 54^. However, cluster 3 cells were not significantly different from the immature cells across any of the measures (Extended Data Fig. 7d-i). Together, this analysis reveals multiple sub-populations of NAc GRPR+ neurons defined by their physiological properties, including putative iSPNs (cluster 1), eSPNs (cluster 0), and immature SPNs (cluster 3).

### GRPR+ neurons in the NAc MSh are more likely to be Fos+ in vivo

Given the high excitability of *Grpr*-eGFP+ cells in our slice recordings, we examined whether they may be more active *in vivo*. To assess this, we performed FISH in naïve wild-type (WT) mice and probed for *Drd2, Grpr*, and *Fos* mRNA. We assessed the percentage of *Drd2*+ cells in the NAc MSh that were *Fos*+ under baseline conditions, using *Fos* as a proxy for neuronal activity. We found that *Drd2*+/*Grpr*+ cells were over twice as likely to be *Fos*+ compared to neighboring *Drd2*+/*Grpr*-cells (18.7% compared to 8.6% *Fos+*) (Fig. 4a,g). Since *Grpr* is expressed in both iSPNs and eSPNs, we repeated the experiment and probed for *Penk* to define iSPNs (Fig. 4b,h) or *Casz1* (Fig. 4c,i) to label eSPNs. In both iSPNs and eSPNs we saw similar results, whereby the *Grpr*-expressing subpopulations were significantly more likely to express *Fos* compared with their *Grpr*-counterparts (Fig. 4h,i).

**Fig. 4:**
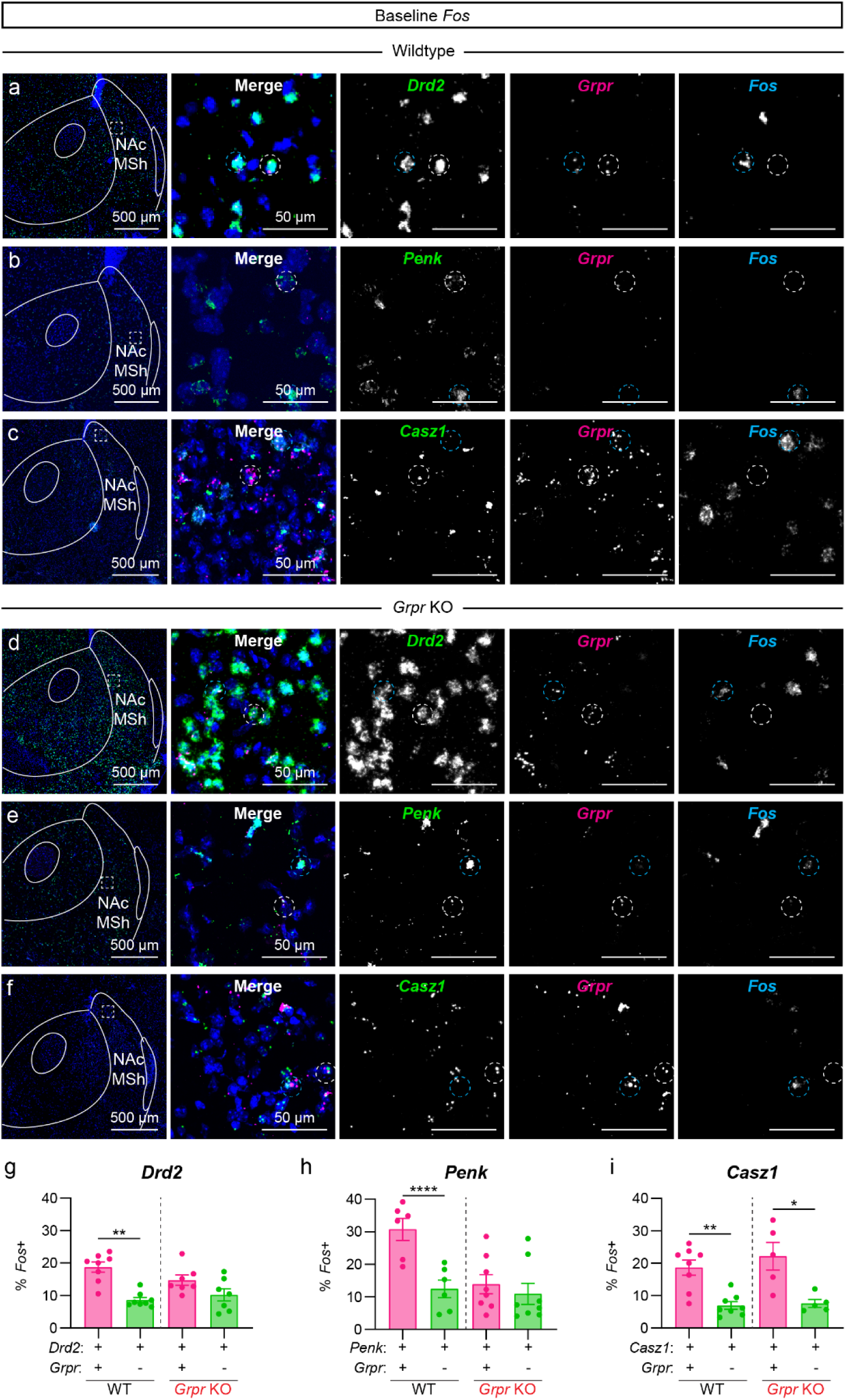
*Grpr+* cells in the NAc MSh have increased *Fos* expression. **(a)** Representative image of the NAc MSh. FISH for *Drd2* in green, *Grpr* in magenta, and *Fos* in cyan. Dashed circles outline sample *Grpr*+ cells, white=*Grpr*+/*Drd2*+/*Fos*-, cyan=*Grpr*+/*Drd2*+/*Fos+*. **(b)** FISH for *Penk* in green, *Grpr* in magenta, and *Fos* in cyan. Dashed circles outline sample *Grpr*+ cells, white=*Grpr*+/*Penk*+/*Fos*-, cyan=*Grpr*+/*Penk*+/*Fos+*. **(c)** FISH for *Casz1* in green, *Grpr* in magenta, and *Fos* in cyan. Dashed circles outline sample *Grpr*+ cells, white=*Grpr*+/*Casz1*+/*Fos*-, cyan=*Grpr*+/*Casz1*+/*Fos+*. **(d-f)** Same as **a**-**c**, but in *Grpr* KO mice. **(g)** Mean ± SEM percentage of *Drd2*+/*Grpr*+ (magenta) vs. *Drd2*+/*Grpr*- (green) cells in the NAc MSh of wild-type (WT) mice (left) and *Grpr* KO mice (right) that are *Fos*+. (WT: n=8 mice; **p=0.0078, Wilcoxon test; KO: n=7 mice; p=0.0781, Wilcoxon test). **(h)** Mean ± SEM percentage of *Penk*+/*Grpr*+ (magenta) vs. *Penk*+/*Grpr*- (green) cells in the NAc MSh of WT mice (left) and *Grpr* KO mice (right) that are *Fos*+. (WT: n=6 mice; ****p<0.0001, Paired t-test; KO: n=8 mice; p=0.0563, Paired t-test). **(i)** Mean ± SEM percentage of *Casz1*+/*Grpr*+ (magenta) vs. *Casz1*+/*Grpr*- (green) cells in the NAc MSh of WT mice (left) and *Grpr* KO mice (right) that are *Fos*+. (WT: n=8 mice; **p=0.0020, Paired t-test; KO: n=5 mice; *p=0.0375, Paired t-test).

To probe whether the increased excitability seen both in slice and *in vivo* was due to GRPR activation or some other mechanism unique to *Grpr*-eGFP+ cells, we generated *Grpr* knockout (KO) mice. To do this, we obtained floxed *Grpr* (*Grpr^fl^*) mice^14^ and bred them to CMV-Cre mice^56^ to induce constitutive deletion of *Grpr* throughout the body. Extended Data Fig. 8 shows PCR genotyping results identifying mice with exon 2 deletion and WT controls. We verified that the *Grpr* FISH probe binding site was not located within exon 2, and thus could be used to identify cells that normally expressed *Grpr*, even if the gene was no longer functional (we term all cells that typically express *Grpr* as “*Grpr*-lineage cells”). We first examined whether there was any difference in the percentage of *Grpr*-lineage cells that were *Fos*+ between WT and *Grpr* KO mice (Extended Data Fig. 8e-g). We found that disrupting *Grpr* expression significantly reduced the percentage of *Grpr*-lineage cells that were *Fos*+ in naïve mice (Extended Data Fig. 8e).

We next repeated the experiments in Fig. 4g-i in the *Grpr* KO mice. In *Grpr* KO mice, there was no longer a significant difference in the percentage of *Fos*+ cells between *Drd2+/Grpr*-lineage cells and *Drd2+/Grpr-* cells (Fig. 4d,g), suggesting that expression of *Grpr* is responsible for their differential activation. Interestingly, when we examined genetically defined iSPNs (*Penk*) and eSPNs (*Casz1*), we saw that while the *Penk*+/*Grpr*-lineage cells were no longer more likely to be *Fos*+ (Fig. 4e,h), the *Casz1*+/*Grpr*-lineage cells retained their high *Fos* expression compared to *Grpr-* cells (Fig. 4f,i). Thus, *Grpr* expression is required for the increased *in vivo* activation of *Grpr*- expressing iSPNs, but not eSPNs.

### Multiple inputs to the NAc MSh express Grp mRNA

While it was previously established that *Grp*+ DA neurons project to the NAc MSh from the VTA^8^, the NAc MSh receives inputs from a number of other brain regions including the hippocampus, amygdala, thalamus, entorhinal cortex, and others^57^. Notably, although GRP and GRPR are relatively sparsely expressed in the brain, several of these brain areas have been reported to express the mRNA for *Grp*^17^. Therefore, we asked whether other inputs into the NAc MSh might also have the potential to co-release GRP. To investigate this, we injected retrobeads into the NAc MSh and performed FISH for *Grp* in the ten primary brain regions that send projections to the NAc MSh^57^ (Fig. 5a,b). We found a high percentage of retrobead+ cells that were also *Grp*+ in the VTA, hippocampus (subiculum and CA1), and basolateral amygdala (BLA) (Fig. 5c,d). We found little to no expression of *Grp* in other regions projecting to the NAc MSh, including the lateral entorhinal cortex, anterior piriform cortex, and thalamus, amongst others (Fig. 5c,d). This suggests that a select group of afferents to the NAc MSh could have the capacity to co-release GRP.

**Fig. 5:**
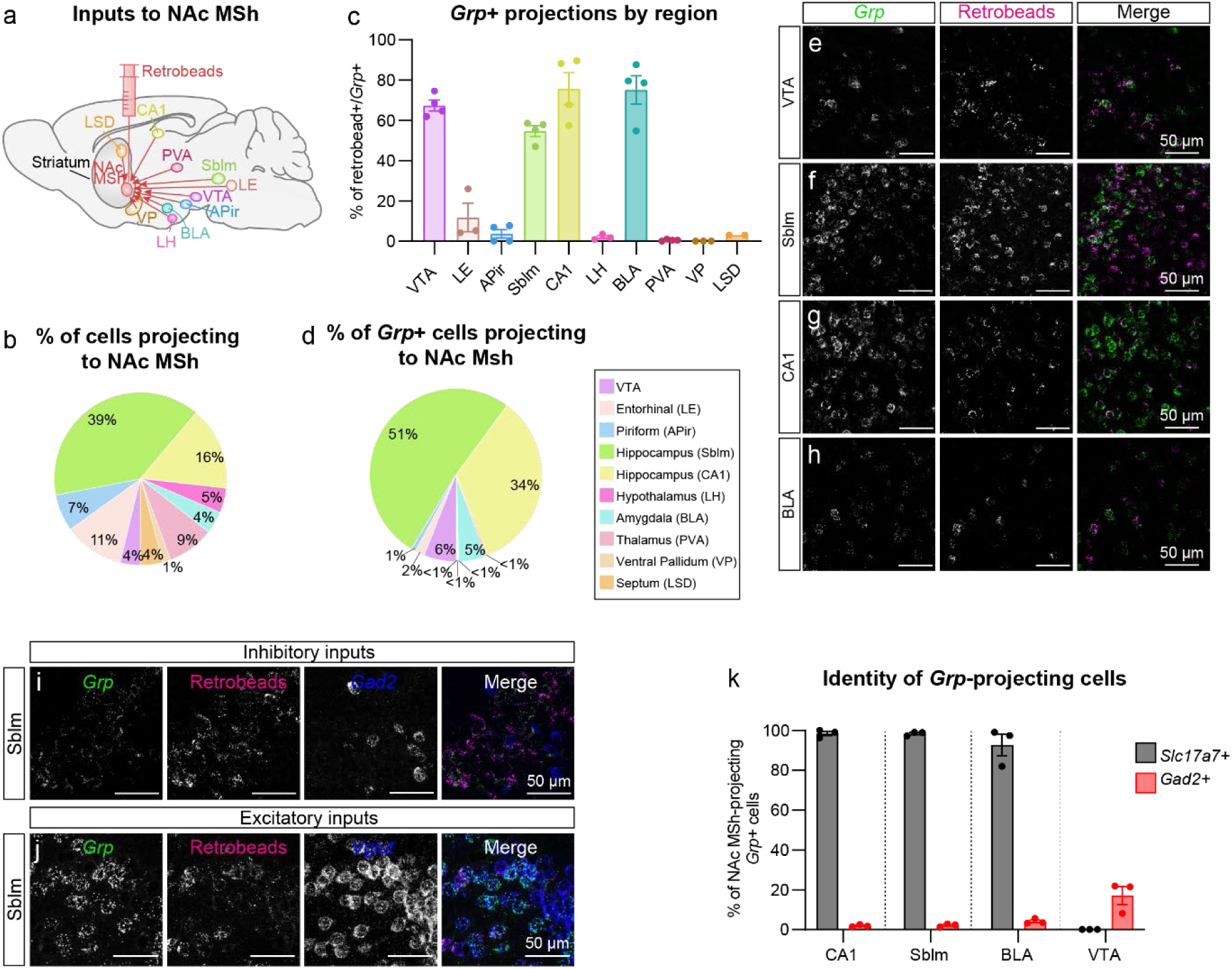
NAc MSh-projecting glutamatergic cells from the hippocampus and amygdala express *Grp*. **(a)** Schematic of retrograde tracing experiment. Red retrobeads were injected into the NAc MSh and the listed projection targets were examined for *Grp* mRNA and retrobead co-localization using FISH. NAc MSh inputs: ventral tegmental area (VTA), lateral entorhinal cortex (LE), anterior piriform (APir), hippocampus subiculum (Sblm), hippocampus CA1 (CA1), lateral hypothalamus (LH), basolateral amygdala (BLA), paraventricular thalamus (PVA), ventral pallidum (VP), and dorsal lateral septum (LSD). **(b)** Pie chart displaying the percentage of total retrobead+ cells coming from each region. Retrobead+ cells were summed across all mice (n=4 injections from 3 mice) and then divided by the number of positive cells in each region. **(d)** Percentage of total retrobead+ cells that were *Grp*+ in each of the examined regions. **(e-h)** Representative FISH images from the indicated brain regions showing *Grp*+ cells in green and retrobead+ cells in magenta; **(e)** VTA, **(f)** subiculum, **(g)** CA1, and **(h)** BLA. **(i**-**j)** Representative FISH images of the subiculum showing *Grp*+ cells in green, retrobead+ cells in magenta, and **(i)** *Gad2*+ or **(j)** *Slc17a7*+ cells in blue. **(k)** Percentage of retrobead+/*Grp*+ cells that were *Gad2+* or *Slc17a7+* for the four main *Grp+* inputs into the NAc MSh (mean ± SEM, dots represent summed value from all slices in that region per 3 unique injection sites from 2 mice).

We summed together all the *Grp*+ inputs quantified and found that the main sources of *Grp* into the NAc MSh were from the subiculum (51%), CA1 (34%), VTA (6%), and BLA (5%) (Fig. 5d-h). We further probed these four regions for *Slc17a7* (excitatory neurons) and *Gad2* (inhibitory neurons) and found that in CA1, the subiculum, and BLA, nearly all the *Grp* inputs were glutamatergic (Fig. 5i-k). This suggests that in addition to the dopaminergic inputs, several different glutamatergic inputs to the NAc MSh may have the ability to use GRP as a neuromodulator.

### GRP depolarizes and increases the excitability of GRPR-expressing neurons

After establishing that GRPR+ cells in the NAc MSh have distinct electrophysiological properties and mapping multiple *Grp*+ inputs to this region, we sought to determine how GRP affects the functional properties of these cells. To do this, we washed-on GRP to NAc MSh slices and measured membrane potential and resistance in *Grpr*-eGFP+ cells. We found that 300 nM GRP caused an approximately 6 mV depolarization in *Grpr*-eGFP+ cells and an average 140% increase in membrane resistance (Fig. 6a,c,d), consistent with the effects of GRP in other cell types^4, 12, 18^. Importantly, these changes were blocked in the presence of the GRPR-specific antagonist, DPDMB (Fig. 6b,e,f). As a control, we performed the same experiment on D2-GFP+ cells in D2-GFP mice^40^, the majority of which are not expected to express GRPR (see Fig. 2n). We did not detect any change in membrane potential or resistance with GRP wash-on (Extended Data Fig. 9), demonstrating that GRP selectively increases the excitability of GRPR-expressing neurons.

**Fig 6:**
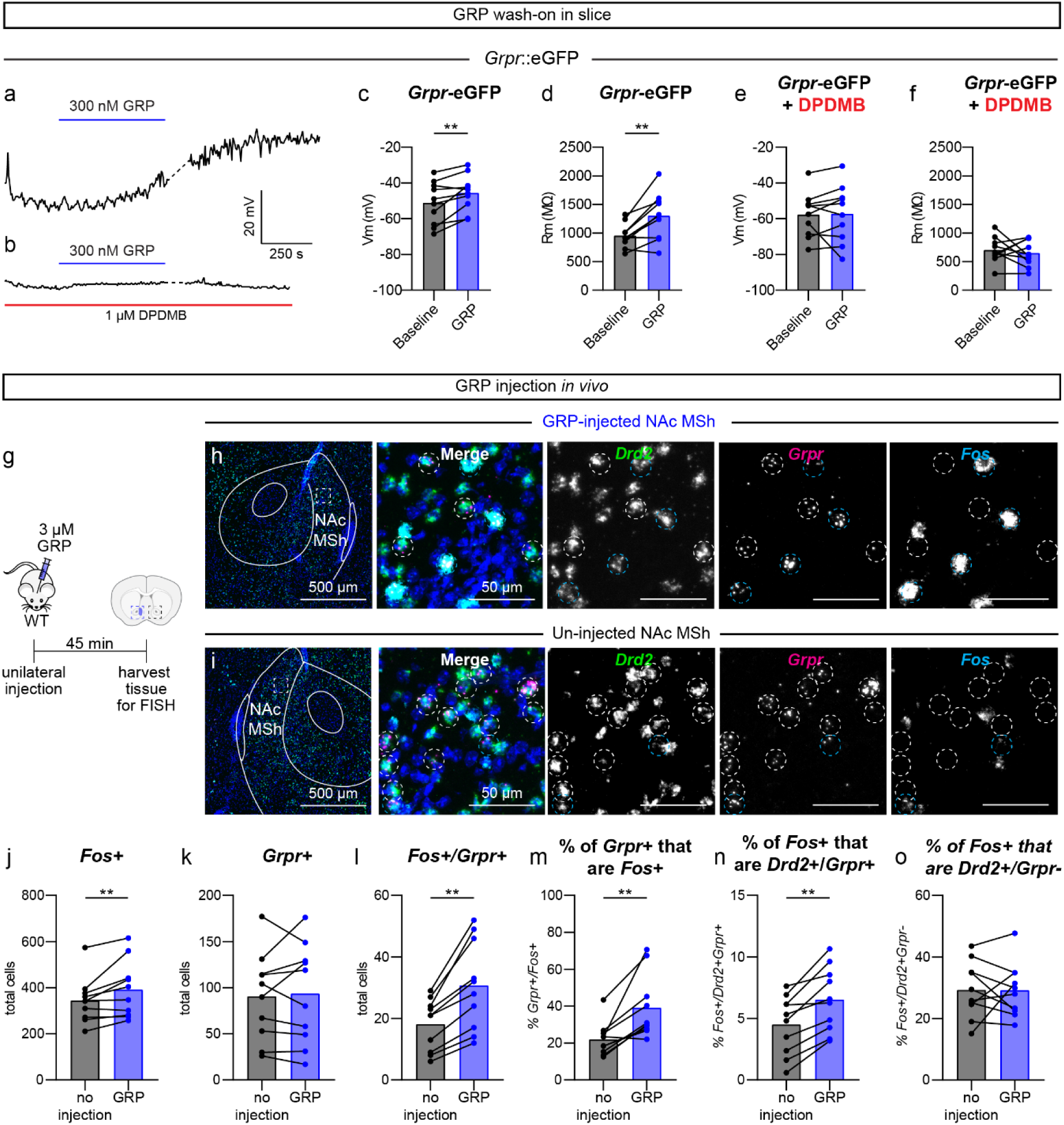
GRP increases the excitability and activation of GRPR-expressing neurons in the NAc MSh. **(a)** Representative trace showing the membrane potential over time of a NAc MSh *Grpr*-eGFP+ cell. The blue bar represents a ten-minute wash-on of 300 nM GRP. **(b)** Representative trace from a NAc MSh *Grpr*-eGFP+ cell in which 300 nM GRP was washed-on for 10 minutes in the presence of 1 μM DPDMB (red bar, entirety of recording). **(c)** Membrane potential of *Grpr*-eGFP+ NAc MSh neurons at baseline (black) and after 300 nM GRP wash-on (blue). Baseline value is the average of the two-minute period prior to GRP wash-on and 300 nM GRP value is the average of the final two minutes of the ten-minute GRP wash-on. (n=10 cells from 7 mice; **p=0.0059, Wilcoxon test). **(d)** Membrane resistance of *Grpr*-eGFP+ NAc MSh neurons at baseline and after GRP wash-on. Baseline value is the average of five sweeps of VC RC check at the beginning of the recording and just following GRP wash-on. (n=10 cells from 7 mice; **p=0.0098, Wilcoxon test). **(e)** Membrane potential of *Grpr*-eGFP+ NAc MSh neurons at baseline and after GRP wash-on in the presence of 1 μM DPDMB; (n=10 cells from 8 mice; p=0.3750, Wilcoxon test). **(f)** Membrane resistance of *Grpr*-eGFP+ NAc MSh neurons at baseline and after GRP wash-on in the presence of DPDMB. (n=10 cells from 8 mice; p=0.6953, Wilcoxon test). **(g)** Schematic of the experiment. 3 μM GRP was injected unilaterally into the NAc MSh of WT mice. 45 minutes post injection, the mice were sacrificed and brains were harvested for FISH. **(h-i)** Representative images of NAc MSh from the **(h)** injected hemisphere and **(i)** un-injected hemisphere. FISH for *Drd2* in green, *Grpr* in magenta, and *Fos* in cyan. Nuclei are labeled with DAPI in blue. Dashed circles outline *Grpr*+ cells, white=*Grpr*+/*Fos*-, cyan=*Grpr*+/*Fos*+. **(j)** Total number of cells in the NAc MSh that were *Fos+* in the un-injected (black) vs GRP-injected hemisphere (blue). (**p=0.0098, Wilcoxon test). **(k)** Total number of cells in the NAc MSh that were *Grpr+* in the un-injected vs GRP-injected hemisphere (p=0.9043, Wilcoxon test). **(l)** Total number of *Grpr*+ cells in the NAc MSh that were *Fos+* in the un-injected vs GRP-injected hemisphere (**p=0.0020, Wilcoxon test). **(m)** Percentage of *Grp*r+ cells in the NAc MSh that were *Fos*+ in the un-injected vs GRP-injected hemisphere (**p=0.0039, Wilcoxon test). **(n)** Percentage of *Fos*+ cells in the NAc MSh that were *Drd2*+ and *Grpr*+ in the un-injected vs GRP-injected hemisphere (**p=0.0020, Wilcoxon test). **(o)** Percentage of *Fos*+ cells in the NAc MSh that were *Drd2*+ and *Grpr*- in the un-injected vs GRP-injected hemisphere (p=0.7695, Wilcoxon test). For panels **j-o**, data were summed across two sections per mouse, n=10 mice.

### GRP injection in vivo increases Fos expression

We next asked whether GRP activates GRPR+ neurons *in vivo* and whether it has broader network effects in the NAc MSh. We injected 3 µM GRP unilaterally into the NAc MSh of anesthetized WT mice and harvested the brains 45 minutes later (Fig. 6g)^18^. We performed FISH and probed for *Drd2*, *Grpr*, and *Fos* mRNA and compared the injected hemisphere to the uninjected hemisphere (Fig. 6h,i). We found an increase in the total number of *Fos*+ cells in the GRP-injected hemisphere compared to the control, indicating an increase in activated cells upon GRP injection (Fig. 6j). The total number of *Grpr+* cells was unchanged, as expected (Fig. 6k). *Fos* was selectively induced in *Grpr*+ cells, as we found an increase in the total number of *Grpr*+ cells that were *Fos*+ as well as an increase in the proportion of *Grpr*+ cells that were *Fos*+ following GRP injection (Fig. 6l,m). Given that *Fos* is a proxy for neuronal activity, these data imply that GRP increases the activity of *Grpr*+ cells in the NAc MSh. We also observed that while the percentage of *Fos*+ cells that were *Drd2+*/*Grpr*+ increased (Fig. 6n), the percentage of *Fos*+ cells that were *Drd2*+/*Grpr*- did not change (Fig. 6o), showing that GRP selectively activated GRPR-expressing cells and did not induce larger network-level activation.

As a control to verify that this effect was induced by GRP and not due to the injection itself, we repeated the above experiment, but injected sterile saline instead of GRP (Extended Data Fig. 10). While we saw a small increase in the total number of *Fos*+ cells in the saline-injected hemisphere, there was no increase in the total number of *Grpr*+ cells or the *Grpr*+ cells that were *Fos*+ (Extended Data Fig. 10a-e). Additionally, we saw no increase in the percentage of *Grpr*+ cells that were *Fos*+ (Extended Data Fig. 10f). Together, these results indicate that while saline injection can acutely increase neuronal activity in the NAc MSh, it does not preferentially activate *Grpr*+ cells.

### Deletion of Grpr in the NAc MSh selectively increases motivation

Given that GRP-GRPR signaling had a strong effect on modulating the excitability of a subset of NAc MSh neurons, we tested whether NAc MSh-specific deletion of *Grpr* affected striatal-dependent behaviors. To generate conditional *Grpr* knock-out (cKO) mice, we injected *Grpr^fl/fl^*or *^fl/y^* mice^14^ bilaterally with an AAV expressing CRE recombinase, to specifically delete *Grpr* in the adult NAc MSh (Fig. 7a,b). A GFP-expressing virus was used as a control. Four weeks post-surgery, *Grpr* cKO mice and control littermates were run through the open field test, accelerating rotarod, progressive ratio (PR) lever-pressing task, and sucrose preference test (Extended Data Fig. 11a) to assess motor and reward-related behaviors.

**Fig. 7:**
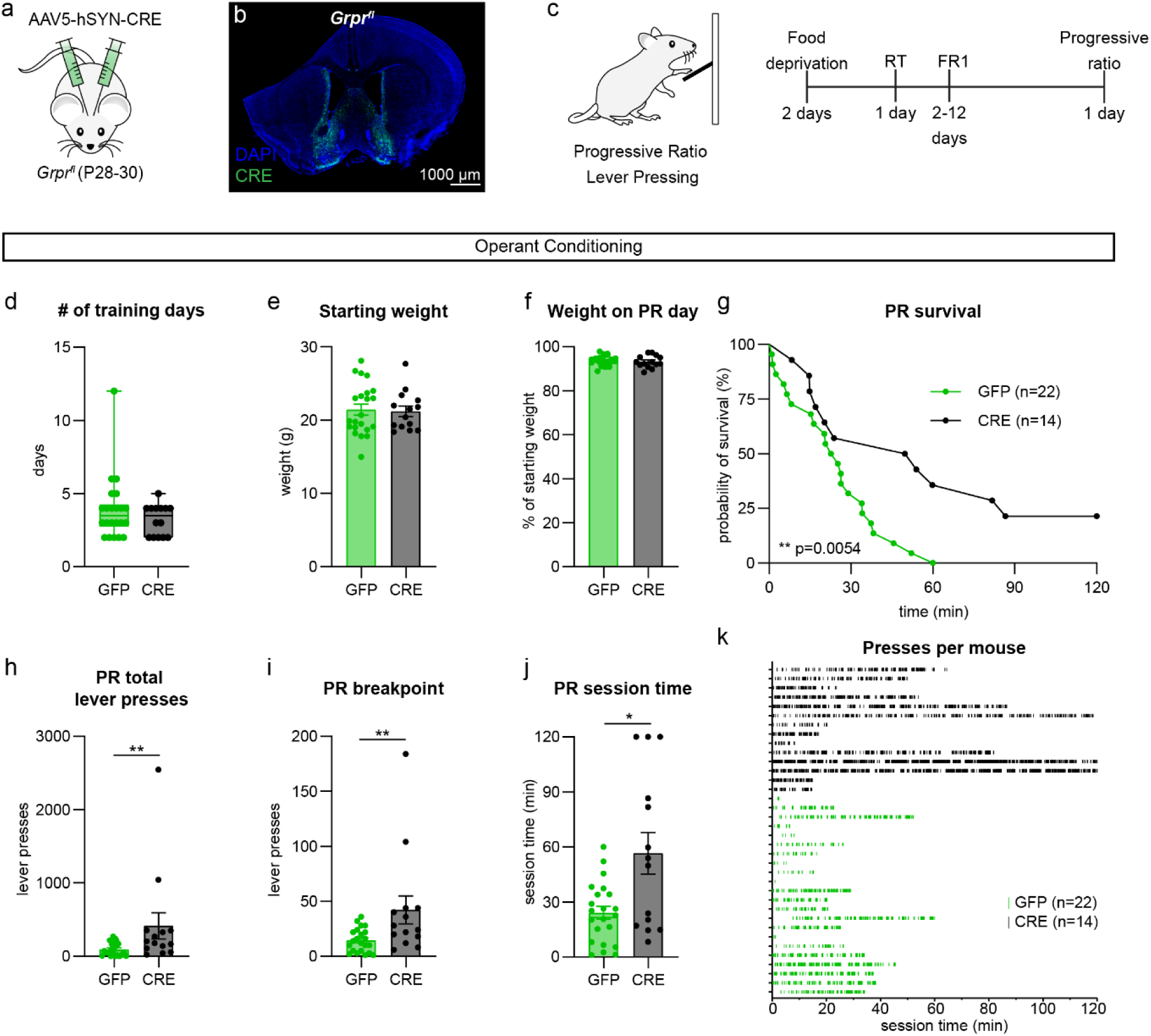
*Grpr* deletion in the NAc MSh enhances motivation in a progressive ratio task. **(a)** Schematic of the experiment. *Grpr^fl/fl^* or *^fl/y^* mice were injected bilaterally at P28-30 with 600 nL of AAV5-hSyn-GFP or AAV5-hSyn-CRE-GFP in the NAc MSh. **(b)** Sample image of injection site in the NAc MSh showing AAV-CRE-GFP in green with nuclei labeled in blue with DAPI. **(c)** Schematic of the progressive ratio (PR) lever pressing task and timeline for operant conditioning. RT = random timing, FR1 = fixed ratio 1. **(d)** Box plots of the number of days it took control (GFP injected, green) and *Grpr* cKO (CRE injected, black) mice to reach criterion on the lever pressing task (boxes represent the interquartile range (25-75%), lines denote the median, whiskers represent mix to max values). Mann-Whitney test, p=0.4522 (n=22 GFP and n=14 CRE mice for all panels). **(e)** Mean ± SEM weight of GFP and CRE mice just prior to starting food deprivation. Mann-Whitney test, p=0.8917. **(f)** Mean ± SEM percentage of starting weight per mouse on the day the mouse completed the PR task. Mann-Whitney test, p=0.4192. **(g)** Survival function of total session time in operant boxes on PR testing day. The session ended when a mouse went three minutes without pressing the lever after pressing the lever at least one time. Mantel-Cox test, **p=0.0054. **(h)** Mean ± SEM total lever presses in the PR session before timeout. Mann-Whitney test, **p=0.0077. **(i)** Mean ± SEM breakpoint in the PR session before timeout. Breakpoint was defined as the number of lever presses for a single reward at which the mouse stopped pressing the lever for more than three minutes. Mann-Whitney test, **p=0.0060. **(j)** Mean ± SEM total session time during the PR task before timeout. Mann-Whitney test, *p=0.0381. **(k)** Raster plot of lever presses per individual mouse during the PR task. Each line represents a single lever press. Each row represents the lever-pressing data of an individual mouse. Green=GFP-injected control mice, Black=CRE injected *Grpr* cKO mice.

The open field assay was used to measure general locomotor behavior and was analyzed using DeepLabCut (Extended Data Fig. 11b)^58^. We found that *Grpr* cKO mice travelled a greater total distance during the one-hour session (Extended Data Fig. 11c), suggesting a mild increase in locomotor activity. We assessed the time spent in the center, as a proxy for avoidance behavior, and number of grooming bouts, as a measure of repetitive behavior, and found no differences (Extended Data Fig. 11d,e). We used Keypoint-MoSeq^59, 60^ to perform unbiased identification of spontaneous behavior syllables used by the mice in the open field. The behavior syllables detected by this approach included movements like forward run, right or left turn, and grooming (Extended Data Fig. 11f,g). We found no significant differences between groups in their usage of any of the top 28 most frequent syllables (Extended Data Fig. 11f). Together, this suggests that disruption of GRPR signaling specifically in the NAc MSh does not have a major impact on general behavior patterns.

Both the DS and VS have been implicated in motor learning^61^. To examine whether *Grpr* deletion impacts motor coordination or motor routine learning, we tested the mice on the accelerating rotarod with three trials per day across four consecutive testing days. We saw no differences in motor coordination or learning between *Grpr* cKO mice and GFP-injected controls (Extended Data Fig. 12a-d).

The NAc is a core component of the mesolimbic reward processing pathway. Alterations in NAc activity can therefore affect the hedonic value of natural rewards^62, 63^. To test whether *Grpr* signaling is involved in this, we compared *Grpr* cKO and control mice in the sucrose preference test (Extended Data Fig. 12e). This test is a measure of hedonia, or the ability to derive pleasure from rewarding stimuli^64–66^. We first verified there was no difference in starting weight between groups (Extended Data Fig. 12f). During the test sessions, we found that *Grpr* cKO and control mice showed a similar preference for the sucrose solution over plain water (Extended Data Fig. 12g). These results suggest that GRPR signaling in the NAc MSh is not required for reward value processing.

In addition to processing reward, the NAc is implicated in the control of motivated behaviors. In particular, modulation of D2 SPNs in the NAc MSh impacts motivation^67, 68^. For example, chemogenetic inhibition of the indirect pathway NAc outputs to the VP is sufficient to enhance motivation^67^, suggesting an inhibitory role for this pathway in motivated behavior. To examine how GRPR signaling impacts motivation, we tested *Grpr* cKO mice on a PR lever pressing task (Fig. 7c). Prior to training, mice were food restricted to 90-95% of their body weight. The mice were trained to press a lever for a food pellet reward in daily sessions with a fixed-ratio of 1 (one press yields one reward, FR1) until they successfully received 20 or more rewards in a one-hour session on two consecutive days. No differences were seen in the number of days it took mice to learn the task (Fig. 7d). There were also no differences in lever-pressing behavior across FR1 training days (Extended Data Fig. 13a,b). Additionally, there was no difference in the starting weight of mice or the percentage of their initial body weight on the PR day between groups (Fig. 7e,f). During the two-hour PR session, mice had to press the lever an increasing number of times to earn food pellet rewards. The PR session was considered over after three minutes had elapsed without a lever press, after the mouse had engaged with the lever at least once. Survival function analysis revealed a significant difference in the number of mice that continued to engage in the task between groups, with more *Grpr* cKO mice demonstrating prolonged pressing (Fig. 7g). Additionally, *Grpr* deletion in the NAc MSh led to a significant increase in the total number of lever presses during the PR session (Fig. 7h), as well as the breakpoint, which is the maximum number of lever presses a mouse is willing to perform for a single pellet reward (Fig. 7i). *Grpr* cKO mice also showed a significant increase in the total PR session time (Fig. 7j,k). These results indicate that disruption of GRP-GRPR signaling in the NAc MSh enhances motivation.

## Discussion

Neuropeptides are potent neuromodulators that regulate neuronal activity and communication in a variety of ways^1, 2^. It has recently become appreciated that many neurons co-express genes for both classical neurotransmitters and one or more neuropeptides^7, 8^. As such, neuropeptides are poised to be key regulators of neural circuit function. Here we identified a novel role for the neuropeptide GRP and its receptor GRPR in mediating the excitability and activity of neurons in the NAc MSh. Specifically, we find that *Grpr*+ neurons in the NAc MSh are a subpopulation of D2R SPNs with distinct electrophysiological properties. The *Grpr*+ cells can be further subdivided into traditional iSPNs as well as eSPNs based on their genetic markers. We mapped the *Grp*-expressing inputs into the NAc MSh and showed that in addition to the previously identified DA neurons from the VTA, the NAc MSh receives significant *Grp+* inputs from glutamatergic neurons in the hippocampus and amygdala. Furthermore, we show both in slice and *in vivo* that the *Grpr*+ neurons in the NAc MSh have high basal excitability and can be further excited by GRP. Finally, we eliminate NAc GRP-GRPR signaling by generating *Grpr* cKO mice and show that this not only reduces the baseline activation of the *Grpr+* neurons, but also leads to increased motivation in a PR task. In this study, we have leveraged gene expression analysis based on FISH and scRNA-seq to identify a previously unknown population of cells in the NAc MSh that plays a functional role in behavior. Additionally, *Grpr* expression provides a unique opportunity to functionally characterize eSPNs for the first time and compare their properties to iSPNs that also express *Grpr*.

### Grpr defines novel subtypes of SPNs within the NAc mSh

The NAc MSh has canonically been thought to be composed of two main types of SPNs that together make up 90-95% of the cells in the region^26^. Traditionally, SPNs were divided into indirect- and direct-pathway neurons based on the type of DA receptor they expressed and their output pathway to other basal ganglia structures^3, 25, 69–71^. dSPNs express D1 receptors, dynorphin, and substance P and have canonically been thought to promote motivated behaviors^34, 72, 73^. In contrast, iSPNs express D2 receptors, adenosine 2A receptors, and enkephalin and generally inhibit motivated behaviors^34, 72, 73^. However, a recent scRNA-seq study identified a third type of SPN, eSPNs^28^. These cells were defined as SPNs based on their expression of classical SPN markers such as *Ppp1r1b*, *Adora2a*, and *Drd1*, but were found to have a unique genetic profile^28^. Since their identification, eSPNs have also been identified in the human striatum, but have not been functionally characterized or compared to their classical SPN counterparts^51^. A recent studying examining a different subpopulation of neurons in the NAc MSh that express *Crh*, also found these cells to be enriched with eSPN markers^55^.

Here we report that *Grpr*-expressing cells in the NAc MSh represent two genetically unique subpopulations, one that co-expresses many canonical iSPN markers including *Drd2*, *Penk*, *Gad2*, and *Ppp1r1b* and another that expresses eSPN markers including *Casz1*, *Th, Otof*, *Cacng5*, and *Pcdh8*. In terms of morphology, all *Grpr*+ cells had spines, and their dendritic morphology was similar to neighboring cells in the NAc MSh, the vast majority of which are expected to be SPNs. Interestingly, when we performed whole-cell current clamp recordings, we found that the *Grpr*-eGFP+ cells were quite different from all other examined cell types in the NAc MSh. While iSPNs in the NAc MSh are considered the most depolarized and excitable SPNs at baseline^3, 74–77^, *Grpr*-eGFP+ cells were both more depolarized and had an approximately 160% greater membrane resistance than *Grpr*-negative *Drd2*+ SPNs. Cluster analysis based on the baseline electrophysiological properties of *Grpr+* cells revealed three subpopulations: one that was similar to traditional iSPNs, one that had the profile of immature iSPNs, and a third group that was distinct from both of these, which we presume represent eSPNs. We found parallel results *in vivo* when we probed for *Fos* mRNA in naïve mice, and found that *Grpr*+ cells were over twice as likely to be *Fos*+ than *Grpr*- iSPNs in the NAc MSh. Interestingly, when we disrupted *Grpr* expression in these cells, *Fos* was no longer preferentially expressed in *Grpr+* compared to *Grpr-* iSPNs, indicating that GRP-GRPR signaling contributes to the baseline excitability of these cells. *Casz1+/Grpr+* cells were also more likely to be *Fos*+ compared to *Casz1+/Grpr-* cells. However, *Fos* expression persisted in the absence of *Grpr* indicating that *Casz1+/Grpr+* cells are more likely to be activated due to reasons independent of the expression of *Grpr*. Together, our detailed analysis reveals the previously unappreciated molecular and functional heterogeneity of cells within the NAc MSh, warranting further investigation into how these different cell types contribute to the computations and output of the NAc.

### Both glutamatergic and dopaminergic inputs into the NAc MSh have Grp

This study was based on prior work showing that a subpopulation of midbrain DA neurons express *Grp*^23, 78, 79^ and project to the NAc MSh^8^. However, many other brain regions are also known to project to the NAc MSh^57^. We found that additional *Grp*-expressing inputs come into the NAc MSh from the hippocampus and amygdala and are glutamatergic. This raises the possibility of co-release of GRP with either DA or glutamate from multiple distinct inputs. Additionally, while neuropeptide signaling has been known to occur in the NAc MSh through opioid receptors, these receptors are all Gi/o-coupled, and the opioids themselves are synthesized within the NAc^3^. In contrast, the *Grp*+ inputs we identified are projections that come into the NAc MSh from other brain regions and excite the target neurons.

GRPR is a Gq-coupled receptor and activation leads to an increase in intracellular calcium^80^. Previous studies examining GRP-GRPR signaling in other brain regions found that GRP excites GRPR-expressing neurons in the auditory cortex, paraventricular thalamus, lateral amygdala, hippocampus and spinal cord^4, 12, 17–21^. In the spinal cord, it was shown that GRP excites neurons via blockade of Kir2 potassium channels^12^. We found that while the *Grpr*+ neurons in the NAc MSh were already spontaneously active at baseline, application of GRP in slice caused further depolarization and increased the membrane resistance by 140%. Importantly, while there are multiple subpopulations of *Grpr*+ cells in the NAc, all cells responded to GRP wash-on. These effects could be blocked by a GRPR-specific antagonist and the presence of the antagonist alone led to a decrease in baseline membrane resistance, raising the possibility that there is basal GRP-GRPR activity. Together these experiments map out a novel mesolimbic GRP-GRPR circuit whereby GRP comes into the NAc MSh from glutamatergic and dopaminergic neurons from the hippocampus, amygdala, and VTA and acutely excites ensembles of GRPR+ cells, which are GABAergic.

### GRP-GRPR signaling affects NAc-associated behaviors

GRP-GRPR signaling in other brains regions is important for several different behaviors^15, 16^. For example, in the amygdala, *Grpr* KO mice showed enhanced long-term fear memories^17^. In the spinal cord, GRP-GRPR signaling is the molecular basis of itch^12–14, 81–83^. More recently, *Grpr* ablation in the auditory cortex decreased auditory fear memories^18^. Given that GRP-GRPR signaling in other brain regions is sufficient to modulate behavior, we generated *Grpr* cKO mice to test whether NAc-associated behaviors were impacted. We did not find major differences in general behavior patterns in the open field, rotarod, or sucrose preference in *Grpr* KO mice indicating that GRPR signaling is not required for these behaviors. We did, however, find that *Grpr* cKO mice showed increased motivation in the PR task. We do not believe this is due to increased lever pressing in general, as there were no significant differences in lever-press behavior during FR1 training. The NAc MSh is an important regulator of motivation and, more specifically, iSPNs have been shown to inhibit motivation^35, 67^. In a prior study, the authors used chemogenetics to inhibit iSPN outputs to the VP in the NAc MSh and found that this increased motivation. This is consistent with studies showing that iSPNs typically inhibit motivated behaviors^3, 29, 34^. Here, by eliminating GRP-GRPR signaling specifically in the NAc MSh, we expected to reduce the excitability of a small subpopulation of iSPNs. Consistent with this, we found that the percentage of *Fos*-positive *Grpr-*lineage iSPNs was reduced in *Grpr* KO mice. The inability to activate *Grpr*-expressing SPNs through GRPR signaling led to increased motivation, consistent with studies broadly manipulating iSPNs in this region. This indicates that GRP-GRPR signaling in the NAc MSh normally constrains motivation and suggests that the relatively small population of GRPR+ NAc neurons can exert significant control over motivated behavior.

### Limitations and future directions

This study identified and characterized previously undescribed subpopulations of SPNs in the NAc MSh that express *Grpr* and *Drd2*. We provide a genetic, morphologic, and electrophysiological characterization of these cells and identify a behavioral role of GRP-GRPR through *Grpr* deletion. These experiments lay the groundwork for further characterization of GRP-GRPR signaling in the striatum and in the brain, in general. We note that although it was beyond the scope of the present study, *Grpr*+ cells also exist in the DS. Given the unique electrophysiological properties of the GRPR+ cells in the NAc MSh, the GRPR+ in the DS warrant their own characterization. Interestingly, there does not appear to be an eSPN population of *Grpr*+ cells in the DS in mice^37^. Additionally, while we identified the *Grp*+ inputs into the NAc MSh and noted that the *Grpr*+ cells themselves were GABAergic, we were unable to address whether the *Grpr*+ cells project to and inhibit the VP. This was largely due to limitations in the available transgenic mouse lines for manipulating *Grpr*-expressing cells. Recent evidence has indicated that unlike the DS, the canonical projection pathways from the NAc are oversimplified and both D1- and D2-SPNs can participate in indirect and direct pathways depending on their downstream target from the VP^35^. Therefore, additional tools are needed to examine the outputs of *Grpr*+ cells. While we perform cellular-level characterization of the *Grpr*-expressing cells and test a subset of behaviors that have been associated with the NAc MSh, future studies could assess a possible role of GRP-GRPR signaling in additional behaviors. One such direction could be utilizing tasks that assess context-induced reinstatement of drug seeking for drugs of abuse. These tests have been associated specifically with ventral subiculum to NAc MSh projections, where we find a large population of *Grp+* inputs to the NAc MSh^84, 85^. Finally, given the Gq-coupling of GRPR, a possible role of GRP in modulating synaptic transmission or plasticity could be examined.

Overall, our study identifies new cell types in the NAc MSh and describes a mesolimbic circuit defined by the expression and functionality of GRP to GRPR signaling.

## Methods

### Mice

Animal experiments were performed in accordance with protocols approved by the University of California, Berkeley, Institutional Animal Care and Use Committee (protocol #: AUP-2016-04-8684-2). Both male and female mice were used across all experiments except for the knock-out FISH experiments where male littermate WT and *Grpr* KO mice were used. Mice were postnatal day P30-P90 unless otherwise stated. The following mouse lines were used in this study: C57BL/6J, JAX strain #000664, *Grpr*::eGFP (GENSAT [MMRRC #036178-UCD])^40^, *Drd2*-eGFP (GENSAT [MMRRC #000230-UNC])^40^, *Drd1*-tdTomato (JAX strain #016204)^42^, CMV-Cre (JAX strain #006054)^56^, *Grpr^fl^* (JAX strain # 033148)^14^, ChAT-IRES-Cre (JAX strain # 06410)^44^, and Ai9 (JAX strain # 07909)^45^. *Grpr* KO mice were generated by breeding CMV-Cre mice to *Grpr^fl^* mice to knock out *Grpr* in all cells.

### Fluorescent *in situ* hybridization

Fluorescent *in situ* hybridization was performed to identify and characterize *Grpr*-expressing cells in the NAc MSh and DS as well as their *Grp*-expressing inputs. Brains were harvested, flash-frozen in OCT mounting medium (Fisher Scientific #23-730-571) on dry ice and stored at -80 °C for up to 6 months. 16 µm-thick coronal sections were collected using a cryostat, mounted directly onto 75x25mm Superfrost® Plus glass slides (VWR #48311-703) and stored at -80°C for up to 1 month. Fluorescent *in situ* hybridization was performed according to protocols provided with the RNAscope® Multiplex Fluorescent Reagent Kit (ACD #320850) and RNAscope® Multiplex Fluorescent Reagent Kit V2 (ACD #323100). *Grpr* mRNA was visualized with a probe in channel 2 (ACD #317871-C2), *Gad2* mRNA in channel 1 (ACD #439371), *Drd2* mRNA in channel 1 (ACD #406501), *Drd1* mRNA in channel 3 (ACD #461901-C3), *Chat* mRNA in channel 1 (ACD #408731), *Ppp1r1b* mRNA in channel 3 (ACD #405901-C3), *Penk* in channel 3 (ACD # 318761-C3), *Casz1* mRNA in channel 3 (ACD #502461-C3), *Fos* mRNA in channel 1 or 3 (ACD #316921 or ACD #316921-C3), *Grp* mRNA in channel 2 (ACD #317861-C2), and *Slc17a7* mRNA in channel 1 (ACD #481851).

Sections were imaged using an Olympus FluoView 3000 confocal microscope equipped with 405, 488, 561, and 640 nm lasers and a motorized stage for tile imaging. Z stack images captured the entire thickness of the section at 1-2.5 μm steps for images taken with 20X (Olympus #UCPLFLN20X) or 10X air objectives (UPLXAPO10X). Additional images were acquired on a Zeiss LSM 710 AxioObserver with Zeiss 10X, 20X, and 63X objectives housed in the Molecular Imaging Center at UC Berkeley. Images were analyzed using FIJI. Cells were considered positive for *Grpr* if they contained two or more fluorescent puncta within the cellular boundary created by DAPI. All images for a single dataset were acquired on the same microscope.

### Immunohistochemistry

Male and female mice (P30-P90) were deeply anesthetized by isoflurane and transcardially perfused with ice cold 1X PBS (∼5–15 mL) followed by 4% paraformaldehyde (PFA) solution (Electron Microscopy Sciences: 15713) in 1X PBS (∼5–15 mL) using a peristaltic pump (Instech). The brains were removed and post-fixed by immersion in 4% PFA in 1X PBS solution overnight at 4°C. Brains were suspended in 30% sucrose in PBS (1X pH 7.4) for cryoprotection. After brains descended to the bottom of the vial (typically 24–48 hr), 40 μm coronal sections of the NAc MSh were cut on a freezing microtome (American Optical AO 860), collected into serial wells, and stored at 4°C in 1X PBS containing 0.02% (w/v) sodium azide (NaN3; Sigma Aldrich).

Free-floating sections were washed with gentle shaking, 3 × 5 min in 1X PBS followed by 1 hr incubation at RT with BlockAid blocking solution (Life Tech: B10710). Primary antibodies were applied at 4°C in 1X PBS containing 0.25% (v/v) Trinton-X-100 (PBS-Tx) overnight to ensure penetration of the antibody throughout the slice. Sections were then washed with cold 1X PBS-Tx 3 × 10 min and incubated for 2 hr at RT with secondary antibodies in PBS-Tx. Sections were washed in cold 1X PBS 3 × 10 min, mounted on SuperFrost slides (VWR: 48311–703), and coverslipped with Prolong Gold antifade mounting media with DAPI (Life Tech: P36935).

We used a primary antibody against GFP (1:5000, Chicken, Abcam: 13970) and an Alexa Fluor 488 goat anti-chicken secondary antibody (1:500, Invitrogen: A-11039).

Images of 40 μm sections processed for immunohistochemistry were acquired using an Olympus FluoView 3000 confocal microscope (described above). Z-stack images captured the entire thickness of the section at 1–2 µm steps for images taken with a 20X air (Olympus #UCPLFLN20X) or a 10X air Olympus #UPLXAPO10X) objective.

### Electrophysiology

Mice (P8-12; P30-90) were perfused transcardially with ice-cold ACSF (pH=7.4) containing (in mM): 127 NaCl, 25 NaHCO3, 1.25 NaH2PO4, 2.5 KCl, 1 MgCl2, 2 CaCl2, and 25 glucose, bubbled continuously with carbogen (95% O2 and 5% CO2). Brains were rapidly removed, and coronal slices (275 μm) were cut on a VT1000S vibratome (Leica) in oxygenated ice-cold choline-based external solution (pH=7.8) containing (in mM): 110 choline chloride, 25 NaHCO3, 1.25 NaHPO4, 2.5 KCl, 7 MgCl2, 0.5 CaCl2, 25 glucose, 11.6 sodium ascorbate, and 3.1 sodium pyruvate. Slices were recovered in ACSF at 34°C for 15 min and then kept at RT before recording.

Recordings were made with a MultiClamp 700B amplifier (Molecular Devices) at RT using 3-5 MOhm glass patch electrodes (Sutter: BF150-86-7.5). Data were acquired using ScanImage software, written and maintained by Dr. Bernardo Sabatini (https://github.com/bernardosabatini/%20SabalabAcq). Traces were analyzed in Igor Pro (Wavemetrics). Recordings with a series resistance > 30 MOhms were rejected.

#### Current-clamp recordings

Current clamp recordings were made using a potassium-based internal solution (pH=7.4) containing (in mM): 135 KMeSO4, 5 KCl, 5 HEPES, 4 Mg-ATP, 0.3 Na-GTP, 10 phosphocreatine, and 1 ETGA. No synaptic blockers were included, and no holding current was applied to the membrane. For intrinsic excitability experiments, positive current steps (1s, +10 to +100 pA) were applied in 10 pA intervals to generate an input-output curve. Membrane potential was averaged over a 300 ms period between 50-350 ms of the 2 s sweep prior to current injection. Membrane resistance was calculated from voltage-clamp RC checks prior to switching to current-clamp. A cell was considered spontaneously active at baseline if it fired at least one action potential during the first two minutes after breaking in without current injection. Action potential half width was calculated as the width at half maximum amplitude of spikes during the first current step that induced spiking. All action potential properties were from the first action potential a given cell fired after current injection.

#### GRP wash-on

Membrane potential changes upon bath application of 300 nM GRP (Anaspec: AS-24214) dissolved in ACSF were monitored in current-clamp mode. 5 s current-clamp recordings were made every 2 s. Prior to GRP wash-on, membrane potential was recorded for 10 minutes at baseline, and membrane potential was averaged over a 300 ms window between 50-350 ms of the 2 s sweep. 300 nM GRP was then bath applied for 10 minutes. At the conclusion of GRP wash-on, we briefly switched back to voltage-clamp to perform 5 RC checks to calculate membrane resistance and confirm that series resistance was stable and <30 MOhms. GRP was then washed-out. The baseline membrane potential was calculated as the average over the two minutes prior to GRP wash-on. The membrane potential following GRP application was calculated as the average over the last two minutes of wash-on. Baseline membrane resistance was calculated from the initial voltage-clamp RC check. Wash-on experiments in the presence of the GRPR antagonist, 1 µM DPDMB (Bachem: H-3042) were performed in the same way, except the slices were pre-incubated in DPDMB for at least 15 minutes prior to recording and DPDMB was present in the ACSF for the duration of the recordings.

### Neurobiotin-filled neuron reconstruction for Sholl analysis

Male and female mice were deeply anesthetized by isoflurane, transcardially perfused with ice cold ACSF using a peristaltic pump (Instech) and decapitated. 275-μm-thick coronal striatal slices were prepared on a vibratome (Leica VT1000 S) in oxygenated ice-cold choline-based external solution (pH 7.8) containing 110 mM choline chloride, 25 mM NaHCO3, 1.25 mM NaHPO4, 2.5 mM KCl, 7 mM MgCl2, 0.5 mM CaCl2, 25 mM glucose, 11.6 mM sodium ascorbate, and 3.1 mM sodium pyruvate. Slices were recovered in ACSF at 34°C for 15 min and then kept at room temperature (RT) for the duration of the recordings. All solutions were continuously bubbled with 95% O2 and 5% CO2. For whole cell recordings, 3–5 mΩ borosilicate glass pipettes (Sutter Instrument: BF150-86-7.5) were filled with a potassium-based internal solution (pH 7.4) containing 135 mM KMeSO4, 5 mM KCl, 5 mM HEPES, 4 mM Mg-ATP, 0.3 mM Na-GTP, 10 mM phosphocreatine, 1 mM EGTA, and 4 mg/mL neurobiotin (Vector laboratories: SP-1120).

275 μm slices containing *Grpr*-eGFP+/- NAc MSh neurons were filled with neurobiotin-containing internal solution (4 mg/mL) during patch-clamp experiments and then fixed in 4% paraformaldehyde solution (Electron Microscopy Sciences: 15713) in 1X PBS for 24–48 hr at 4°C. With continuous gentle shaking, slices were washed in 1X PBS 3 × 5 min and incubated with BlockAid blocking solution (Life Tech: B10710) for 1.5 hr at RT. Streptavidin Alexa Fluor 633 conjugate and Streptavidin Alexa Fluor 488 (1:2000 (633) 1:1000 (488), Invitrogen: S21375 (633) S32354 (488)) was applied for 2 hr in 1X PBS containing 0.50% (v/v) Triton-X-100 (PBS-Tx). Slices were then washed in 1X PBS 3 × 10 min, mounted on SuperFrost slides (VWR: 48311– 703) with the cell-containing side of the slice facing up, and coverslipped with Prolong Gold antifade mounting media with DAPI (Life Tech: P36935).

Filled cells in the mounted sections were imaged on an Olympus FluoView 3000 confocal microscope at 20X magnification with 2X digital zoom for a final magnification of 40X. 1 μm steps were used to acquire a z-stack spanning the entirety of the neurobiotin-filled cell body and dendritic arbors. 3D reconstructions of the cells were done using IMARIS 9.2.1 software (Bitplane) with automated filament tracing and manual editing. The mask generated by the automated filament tracing algorithm was continuously cross-referenced with the original z-stack image to ensure accuracy. Spurious segments created by the automated filament tracer due to background noise were removed, while processes with incomplete reconstruction were manually edited to incorporate missing segments.

### Sparse viral injections for spine density quantification

Neonatal (P0) or juvenile (P28-30) *Grpr*::eGFP mice were injected bilaterally with 250 nL (neonates) or 100 nL (juveniles) AAV5-hSyn-mCherry (Addgene: 114422-AAV5) diluted 1:200 to label neurons in the NAc MSh. Neonatal mice were cryoanesthetized and injected bilaterally at 200 nL/min. Injections were targeted to the NAc MSh, with coordinates approximately 0.75 mm lateral to midline, 1.0 mm posterior to bregma, and 3.0 mm ventral to the head surface. For juvenile injections, mice were briefly anesthetized with 3% isoflurane (Piramal Healthcare: PIR001710) and oxygen. Mice were mounted on a stereotaxic frame (Kopf instruments Model 940) with stabilizing ear cups and a nose cone delivering constant 1.5% isoflurane in medical oxygen. Viruses were injected using a pulled glass capillary (Drummond Scientific Company: 5-000-2005) at a rate of 100 nL/minute. Following injection, the capillary remained in place for 1 minute per every minute spent injecting to allow the tissue to recover and prevent virus backflow up the injection tract upon retraction. Coordinates from bregma for juvenile injections were as follows: M/L +/- 1.00 mm, A/P +1.90 mm, D/V -4.50 mm.

All mice were perfused at P56. Mice were perfused and brains were post-fixed with 4% paraformaldehyde overnight, then sectioned at 80 μm. Sections were blocked for 1 hr at RT in BlockAid (ThermoFisher: B10710) and incubated for 16 hr at 4°C with antibodies against RFP (1:500, rabbit, Rockland (VWR): RL600-401-379) and GFP (1:500, chicken, Abcam: 13970). Sections were washed 3 × 10 min in PBS-Tx and incubated for 2 hr at RT with Alexa Fluor 488 and Alexa Fluor 546 secondary antibodies (1:500, Invitrogen: A-11039 and A-11001). Sections were washed 3 × 10 min in 1X PBS and mounted onto slides using Prolong Gold antifade mounting media with DAPI (Life Tech: P36935).

Dendrites were imaged on an Olympus FluoView 3000 scanning confocal microscope at 60X magnification with 2.5X digital zoom for a final magnification of 150X. Images were deconvoluted in the CellSens4 software using a built-in advanced maximum likelihood algorithm. Dendritic spine reconstructions were generated using the filament tracer feature in Imaris 9.3.1. 1-9 dendrites were reconstructed per cell, with the detection parameters for thinnest spine diameter set to 1.5 µm and the maximum spine length set to 3.5 µm. After automatic detection of spines, the digital reconstruction was compared to the original z-stack image to ensure accuracy. Detected spines were manually reviewed by the experimenter and added or removed as needed. Spine density of each dendrite was calculated by dividing the number of spines on that dendrite by the total dendrite length. The values of all dendrites quantified for a given neuron were averaged and the average spine density per dendrite is reported.

### Single-cell RNAseq analysis

scRNAseq data was downloaded from the Allen Brain Cell Atlas (Mouse:^37^, Human:^51^). Expression matrices and metadata for relevant brain regions were extracted, processed, and analyzed using Seurat v.5.1.0^86^. For mouse, data corresponding to the ventral (STRv) and dorsal (STRd) striatum were selected. For human, data corresponding to the dissection region “Basal nuclei (BJ) - Nucleus Accumbens - NAC” were selected. Striatum datasets were processed using the “Standard Seurat Workflow” (https://satijalab.org/seurat/articles/essential_commands.html). Subsetted Seurat objects were re-processed using this workflow before additional analysis and plotting. To produce GRPR-only datasets, any cell expressing non-zero levels of *Grpr*/*GRPR* were subsetted from the overall Seurat objects, and re-processed using the Standard Seurat Workflow. Data for gene expression violin plots was extracted from the log-normalized RNA assay.

### Passive properties clustering analysis

Passive properties were directly visualized using the pairplot function in the Seaborn Python package^87^. Passive properties were analyzed using the UMAP-learn^88^ and Scikit-learn^89^ Python packages. Briefly, data was pre-processed using the StandardScaler function, and UMAP embeddings were calculated using the following parameters: (n_neighbors=10, min_dist=0, metric=’euclidean’). Clustering was performed on the scaled dataset using the K-means approach with an n_clusters value of 4.

### Intracranial injections

#### Retrobead injections

P50-90 WT male and female mice were used for retrograde labeling experiments. Red retrobeads IX (Lumafluor: 78R180) were diluted 1:3 in sterile saline. 350 nL diluted beads were unilaterally injected into the NAc MSh using a pulled glass pipette. The following coordinates from bregma were used: M/L + 0.65 mm, A/P +1.90 mm, D/V -4.50 mm. To allow for sufficient labeling, mice were sacrificed 14 days post injection. Brains were harvested, flash-frozen in OCT mounting medium (Fisher Scientific: 23-730-571) on dry ice and stored at -80°C for up to 6 months. 16 µm-thick coronal sections were collected using a cryostat, mounted directly onto 75x25mm Superfrost® Plus glass slides (VWR: 48311-703) and stored at -80°C for up to 1 month. Prior to sectioning the posterior part of the brain, the injection site was validated to ensure that retrobeads were constrained to the NAc MSh. After confirming, the rest of the brain was sectioned and fluorescent *in situ* hybridization was performed on all sections containing brain regions with retrobead+ cells.

#### GRP injections

Male and female mice (P30-P90) underwent stereotaxic surgery to inject GRP into the NAc MSh. 800 nL of 3 µM GRP (dissolved in sterile saline) or sterile saline were injected unilaterally into the left NAc MSh. The stereotaxic coordinates used were: + 0.65 mm, A/P +1.90 mm, D/V -4.50 mm. 45 minutes after the end of the injection, mice were sacrificed, and their brains harvested and flash frozen for fluorescent *in situ* hybridization.

#### AAV-CRE and GFP injections

Male and female mice (P28-30) underwent stereotaxic surgery to inject pENN.AAV.hSyn.HI.eGFP-Cre.WPRE.SV40 (AAV5) or pAAV-hSyn-EGFP (AAV5) into the NAc MSh. 600 nL of 1:3 diluted CRE or GFP virus in sterile saline were injected bilaterally into the NAc MSh. The stereotaxic coordinates used were: +/- 0.85 mm, A/P +1.70 mm, D/V -4.50 mm. Mice were allowed to recover and viruses were given four weeks to express before mice underwent behavioral testing. At the end of behavioral testing, mice were sacrificed and the tissue was sliced to confirm proper viral targeting.

### Genotyping to validate global KO mice

Multiple PCRs were used as a genotyping strategy to confirm the genotype of *Grpr* KO and WT control mice. The strategy involved primer pairs for the presence of the floxed or WT allele, the LoxP scar, and an internal region of *Grpr* exon 2. The following primer sets were used: *Grpr* floxed (WT-forward: 5’ GGAGAGAGGTATAGAGGGGC 3’, WT-R: 5’ ACATGTAGGTTGCAGGGGAT 3’, MUT-F 5’ GGGTTATTGTCTCATGAGCGG 3’), LoxP scar (Forward 5’ AGACAGCTCTATCACGGTCC 3’, Reverse 5’ GAGTTCAGTTCCCAGCAACC 3’), *Grpr* Exon 2 (Forward 5’ TTTCTGACCTCCACCCCTTC 3’, Reverse 5’ CACGGGAAGATTGTAGGCAC 3’).

With this strategy, we expected the following bands for *Grpr* KO mice: *Grpr* floxed (no band), LoxP scar (298 bp), *Grpr* Exon 2 (no band) and WT mice: *Grpr* floxed (573 bp WT band), LoxP scar (no band), *Grpr* Exon 2 (206 bp).

### Behavioral experiments

Behavior studies were carried out in the dark phase of the light cycle under red lights (open field) or white lights (rotarod and operant conditioning). Sucrose preference was carried out in the room where the mice are normally housed. Mice were habituated to the behavior testing room for at least 30 minutes prior to testing, and at least 24 hours elapsed between sessions. All behavioral equipment was cleaned between each trial with 70% ethanol. Equipment was rinsed with diluted soap and water at the end of each day. Behavioral experiments included both male and female mice and began four weeks after the stereotaxic injections at P54-P60. Male mice were trained or tested before female mice each day. Experimenters were blinded to the genotype during behavioral testing.

#### Open field

Exploratory behavior in a novel environment and general locomotor activity were assessed by a 60 min session in an open field chamber (40 cm L x 40 cm W x 34 cm H) made of transparent plexiglass. The mouse was placed in the bottom right hand corner of the arena and behavior was recorded using an overhead monochrome camera (FLIR Grasshopper 3: GS3-U3-41C6NIR-C) with a 16 mm wide angle lens (Kowa: LM16HC) placed on top of the arena from a height of 50 cm. Data was analyzed using DeepLabCut (DLC)^58, 90^ and Keypoint-MoSeq^59, 60^.

#### Processing and analysis of open field data

To extract the body part (keypoint) coordinates from the video recordings, we used DLC 2.3.4^58,90^. Fourteen body parts including nose, head, left ear, right ear, left forelimb, right forelimb, spine 1, spine 2, spine 3, left hindlimb, right hindlimb, tail 1, tail 2, and tail 3 were manually labelled on a small subset of the video frames. A DLC model was then trained using the annotated frames to label these 14 body parts for all videos recorded. The distance travelled and center entries were calculated with the coordinate of body part tail 1.

Discrete behavior syllables were extracted with Keypoint-MoSeq 0.4.4^60^. Syllable usage and transition data were obtained using built-in functions in the Keypoint-MoSeq package. Decoding analysis was done with customized Python 3.9 script. Code will be made available in Github prior to publication.

#### Rotarod

The accelerating rotarod test was used to examine motor learning. Mice were tested on a rotarod apparatus (Ugo Basile: 47650) for four consecutive days. Three trials were completed per day with a 5 min break between trials. The rotarod was accelerated from 5-40 revolutions per minute (rpm) over 300 s for trials 1-6 (days 1 and 2), and from 10-80 rpm over 300 s for trials 7-12 (days 3 and 4). On the first testing day, mice were acclimated to the apparatus by being placed on the rotarod rotating at a constant 5 rpm for 60 s and returned to their home cage for 5 minutes prior to starting trial 1. Latency to fall, or to rotate off the top of the rotarod barrel, was measured by the rotarod stop-trigger timer.

#### Operant conditioning

Motivated behavior was assessed in an operant conditioning lever pressing task using a food pellet reinforcer. Animals were food restricted to ∼90-95% of their body weight before commencing training in an operant conditioning chamber (Med Associates: ENV-307A). The chambers contained a single retractable lever on the left side of the food receptacle and a house light on the opposite end of the chamber. Each session began with illumination of the house light and the presentation of the left lever. The session ended with retraction of the lever and with the house light turning off. The lever remained present throughout the session. Mice were weighed before each session and habituated to the experiment room for at least 30 minutes prior to testing.

During the first training session, following two days of food restriction, a food pellet (14 mg regular “chow” pellet, Bio-Serve: F05684) was delivered on a random time schedule (RT), with a reinforcer delivered on average every 60 seconds for a total of 15 minutes with no levers present. On subsequent days, mice underwent 60-minute fixed ratio (FR1) sessions where mice were presented with only the left lever. Each lever press was rewarded with a single pellet reward. During initial FR1 sessions, crushed pellets were placed on the lever at the beginning of the task and each time 10 minutes elapsed without a lever press to prime the mice to press the lever. Mice underwent daily FR1 sessions until they had two consecutive sessions with 20 or more lever presses. After earning 20 rewards the first time, the levers were no longer primed with crushed pellets. The mice then moved on to the progressive ratio (PR) session the following day. The PR session ended when a mouse failed to press the lever for a period of at least three minutes, after initiating the lever at least once, or after two hours had elapsed, whichever came first. The number of lever presses required for a reward increased with each reward. The number of lever presses to earn a reward were as follows: 1, 2, 3, 4, 5, 6, 7, 8, 10, 12, 14, 16, 18, 20, 22, 24, 28, 32, 36, 40, 44, 48, 52, 56, 64, 72, 80, 88, 96, 104, 112, 120, 128, 136.

#### Sucrose preference test

*Grpr* cKO and GFP-injected control mice were assessed for normal hedonic behavior by using the sucrose preference test. Three days prior to the sucrose preference test, the water source in the home cage was switched from a Hydro Pac (plain water) to a water bottle containing the contents of the Hydro Pac. After three days of acclimating to the water bottles, a second bottle was added to the home cage containing 5% sucrose in Hydro Pac water. Mice had access to both bottles for 24 hours. After 24 hours, mice were individually housed, again with two bottles, one with 5% sucrose and the other with standard water. Mice were kept singly housed for another 24

hours. The volume of water and sucrose consumed during this 24-hour period was measured. Sucrose preference was calculated by taking the total volume of sucrose solution consumed and dividing it by the total liquid consumed in the 24-hour period. To avoid any confounds due to side-specific preferences, the bottles’ location was randomly assigned (front or back) for each mouse. All experiments were conducted on weekends, ensuring a quiet environment for the tested animals.

### Quantification and statistical analysis

GraphPad Prism 10 was used to perform statistical analyses. All datasets were first analyzed using D’Agostino and Pearson normality test, and then parametric or non-parametric two-tailed statistical tests were employed accordingly to determine significance. If the variances between two groups were significantly different, a Welch’s correction was applied. Significance was set as *p<0.05, **p<0.01, ***p<0.001, and ****p<0.0001. All p values were corrected for multiple comparisons. Statistical details including sample sizes for each experiment are reported in the figure legends.

## Acknowledgments

This work was supported by a Chan Zuckerberg Biohub Investigator grant to H.S.B. We thank Dr. Daniel Kramer for his contribution to the early phases of the project. We thank Dr. Hendrik Wildner and Dr. Hanns Ulrich Zeilhofer for sending us tissue samples for mouse validation. We thank Kamran Ahmed for help with schematics for figures. We thank Dr. Dirk Hockemeyer for help designing a genotyping strategy for the *Grpr* KO mice. Select confocal imaging experiments were conducted at the CRL Molecular Imaging Center, RRiD:SCR_017852, supported by the Gordon and Betty Moore Foundation. We thank Holly Aaron and Feather Ives for their microscopy advice and support. We thank Mahmoud Farhan for help with mouse colony maintenance and genotyping.

## Author contributions

Conceptualization – E.E.A. and H.S.B.; Formal analysis – E.E.A., T.L.L., and H.W.; Funding acquisition – H.S.B.; Investigation – E.E.A., T.L.L, H.W., A.A.S., and E.M.T.; Methodology – E.E.A., T.L.L., H.W., and E.M.T.; Project administration – E.E.A. and H.S.B., Software – T.L.L. and H.W.; Supervision – H.S.B.; Visualization – E.E.A.; Writing – original draft – E.E.A.; Writing – review & editing – E.E.A., T.L.L., H.W., A.A.S., E.M.T., and H.S.B.

---

**Extended Data Fig. 1:**
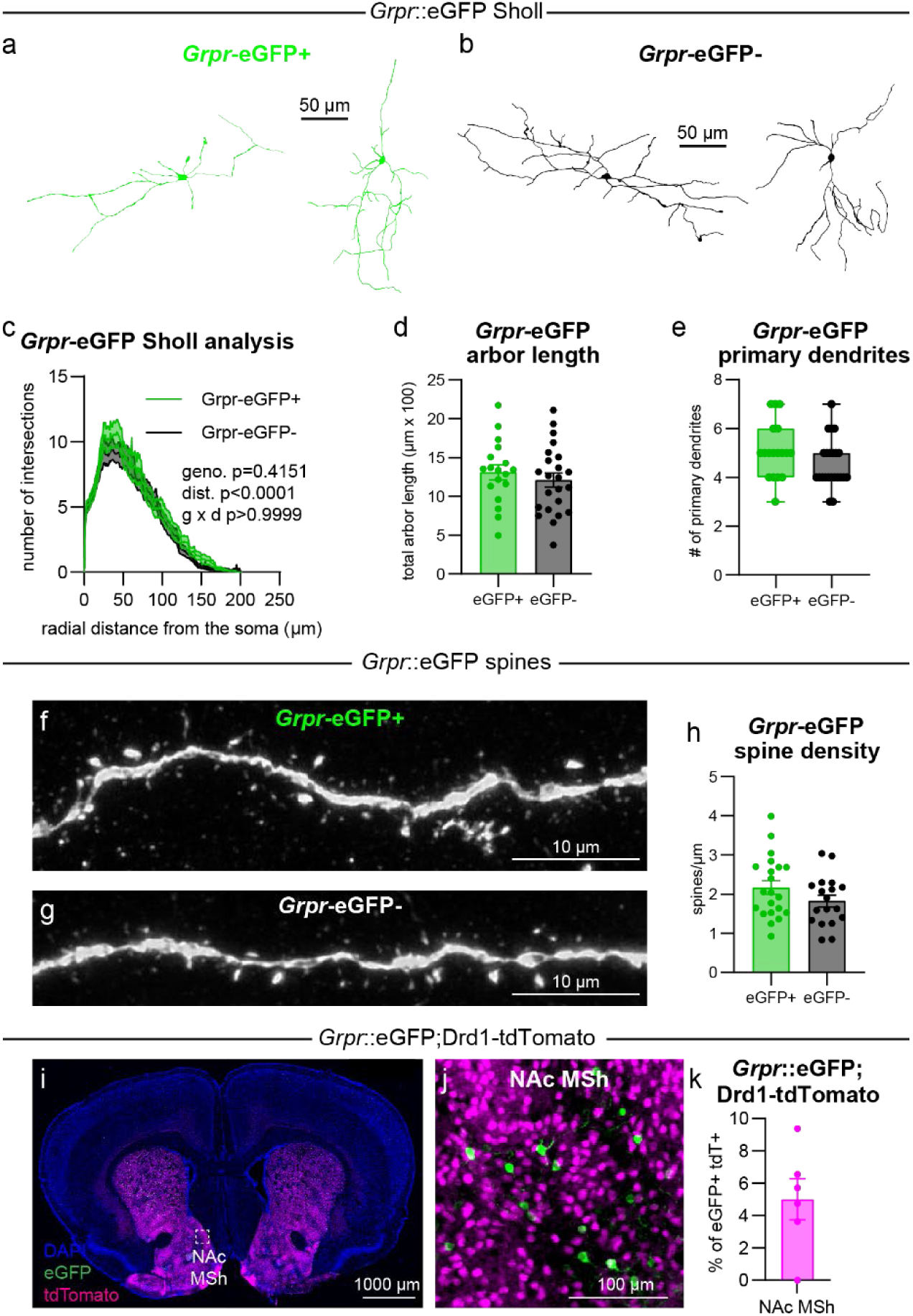
GRPR-expressing neurons in the NAc MSh are morphologically similar to neighboring neurons. **(a-b)** Reconstructions of the dendrites and cell body of NAc MSh **(a)** *Grpr*-eGFP+ and **(b)** *Grpr*- eGFP-neurons. **(c)** Sholl analysis of NAc MSh neuron dendritic arbors. Dark colored lines are the mean, lighter color shading represents the SEM. *Grpr*-eGFP+ in green: n=18 neurons from 8 mice, *Grpr*-eGFP- in black: n=23 neurons from 14 mice. Two-way ANOVA p-values are shown. **(d)** Mean ± SEM total dendritic arbor length per cell. n is the same as for panel **c** (p=0.4452, Welch’s two-tailed t-test). **(e)** Box plots of the number of primary dendrites per neuron for each genotype (boxes represent the interquartile range (25-75%), lines denote the median, whiskers represent mix to max values). n is the same as for panel **c** (p=0.1151, Welch’s two-tailed t-test). **(f**-**g)** Representative images of mCherry-expressing dendrites from a **(f)** *Grpr*-eGFP+ and **(g)** *Grpr*-eGFP-neuron in the NAc MSh. **(h)** Mean ± SEM spine density in *Grpr*-eGFP+ neurons vs *Grpr*-eGFP-neurons. *Grpr*-eGFP+ in green: n=21 neurons from 11 mice, *Grpr*-eGFP- in black: n=18 neurons from 9 mice; p=0.1357, Welch’s two-tailed t-test. For panels **d**, **e**, and **h**, dots represent values from individual neurons. **(i)** Representative coronal brain section (A/P +1.70) from a *Grpr*::eGFP;Drd1-tdTomato mouse. **(j)** Zoom-in of the NAc MSh from panel **i**. **(k)** Mean ± SEM percentage of eGFP+ cells that are tdTomato+ in *Grpr*::eGFP;Drd1-tdTomato mice in the NAc MSh (n=6 mice). Related to Figure 1.

**Extended Data Fig. 2:**
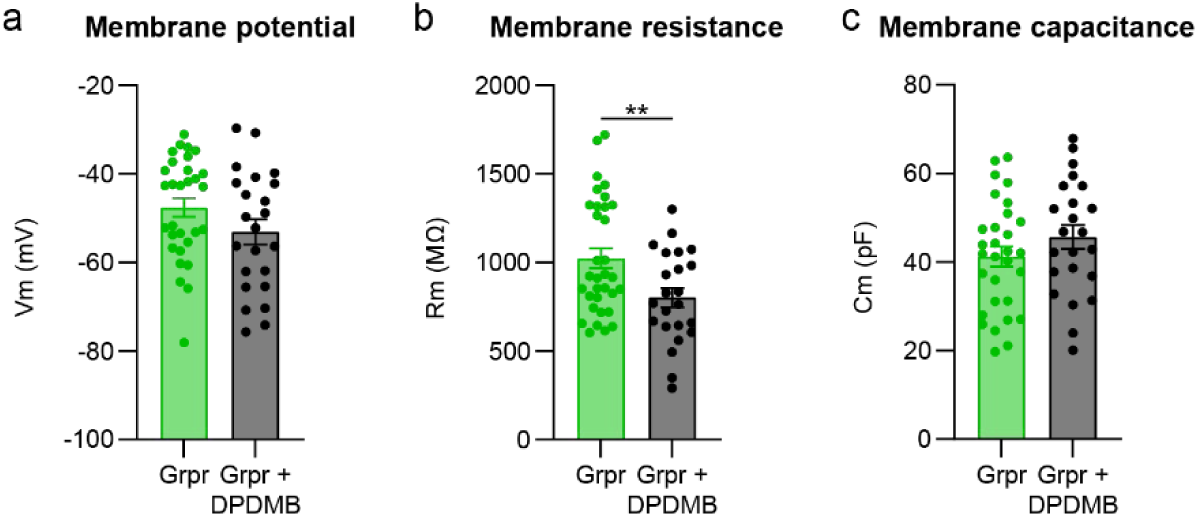
DPDMB decreases the membrane resistance of GRPR-expressing neurons in the NAc MSh. **(a)** Mean ± SEM membrane potential of *Grpr*-eGFP+ NAc MSh neurons. *Grpr*-eGFP+ (no DPDMB) in green: n=30 cells from 15 mice and *Grpr*-eGFP+ (1 μM DPDMB) in black n=23 cells from 9 mice for all panels (Welch’s two-tailed t-test, p=0.1288). **(b)** Mean ± SEM membrane resistance of *Grpr*-eGFP+ NAc MSh neurons with (black) and without (green) DPDMB (Welch’s two-tailed t-test, **p=0.0066). **(c)** Mean ± SEM membrane capacitance of *Grpr*-eGFP+ NAc MSh neurons with (black) and without (green) DPDMB (Welch’s two-tailed t-test, p=0.2199). Related to Figure 1.

**Extended Data Fig. 3:**
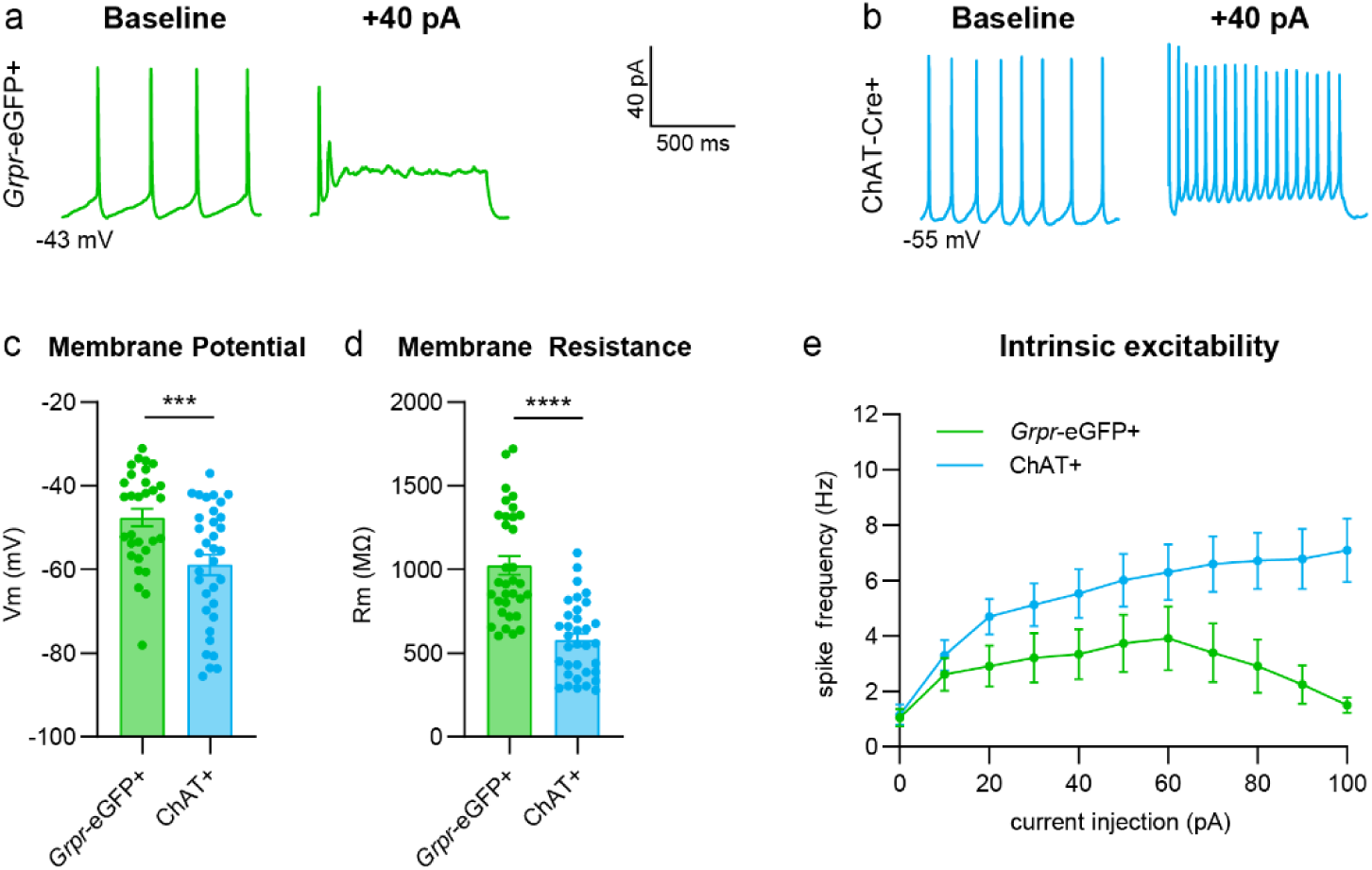
GRPR-expressing neurons in the NAc MSh have distinct electrophysiological properties from cholinergic interneurons. **(a-b)** Representative traces from a **(a)** *Grpr-*eGFP+ and **(b)** *Chat*-Cre+ neuron in the NAc MSh recorded at baseline (left) and following +40 pA current injection (right). **(c)** Mean ± SEM membrane potential of NAc MSh neurons. *Grpr*-eGFP+ in green: n=30 cells from 15 mice and *Chat*-Cre+ in light blue: n=34 cells from 7 mice (Welch’s two-tailed t-test, ***p=0.0009). **(d)** Mean ± SEM membrane resistance of NAc MSh neurons. *Grpr*-eGFP+ in green: n=33 cells from 15 mice and *Chat*-Cre+ in light blue: n=34 cells from 7 mice. (Welch’s two-tailed t-test, ****p<0.0001). **(e)** Input-output curves showing the mean ± SEM firing frequency of NAc MSh neurons in response to positive current steps of increasing amplitude. *Grpr*-eGFP+ in green: n=23 cells from 10 mice and *Chat*-Cre+ in light blue: n=34 cells from 7 mice. (Two-way repeated measures ANOVA, current injection x cell type ***p=0.0008, current injection **p=0.0010, cell type **p=0.0079). Data and traces for *Grpr*-eGFP+ cells are duplicated from Fig. 1 for comparison. Related to Figure 1.

**Extended Data Fig. 4:**
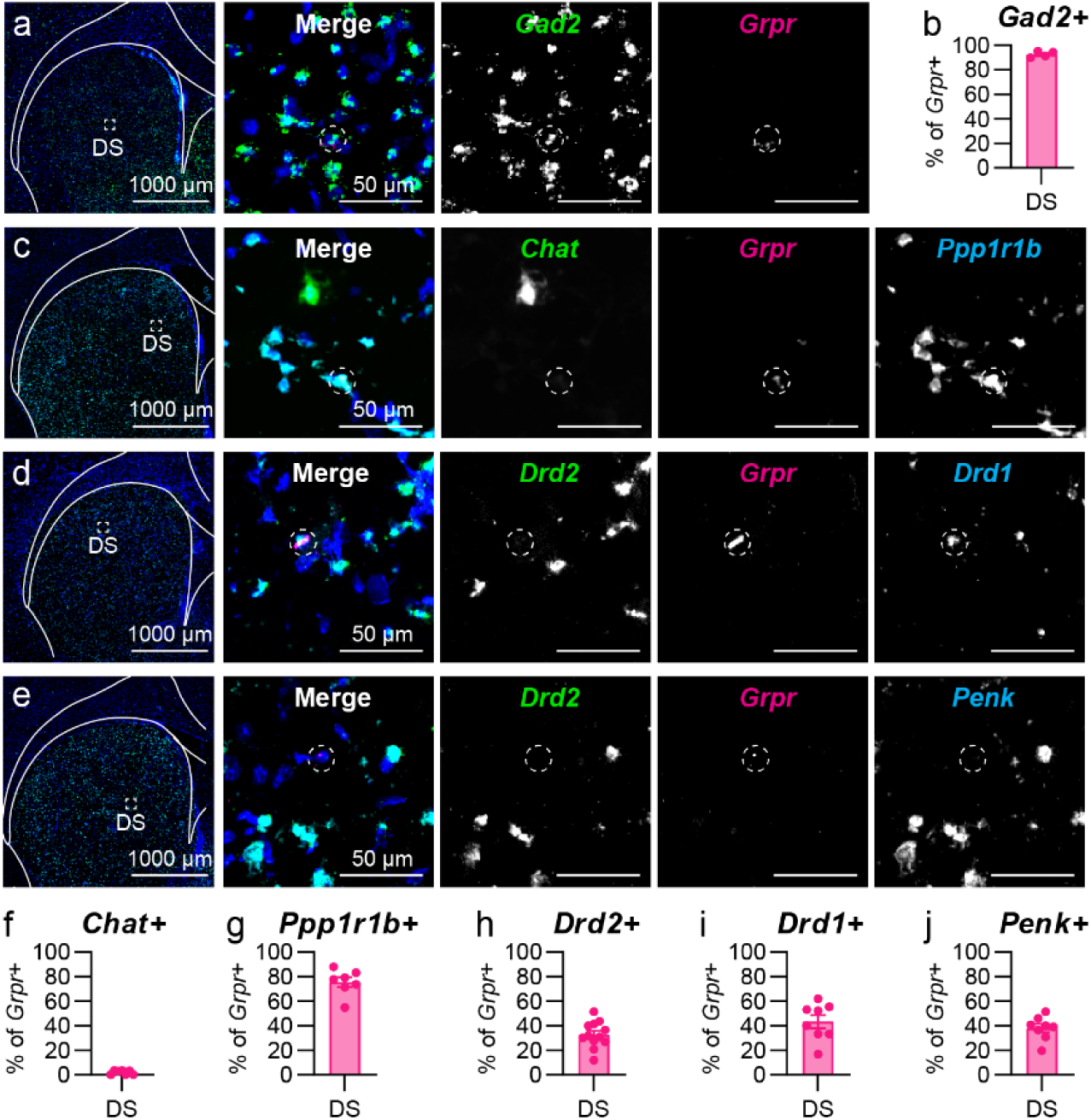
*Grpr* is expressed in SPNs in the dorsal striatum. **(a)** Representative image of the DS. FISH for *Gad2* in green and *Grpr* in magenta. Nuclei are labeled with DAPI in blue. Right panels show zoomed-in images of the boxed region. Dashed circles outline *Grpr+* cells. **(b)** Mean ± SEM percentage of *Grpr*+ cells in the DS that are *Gad2+* in WT mice. (n=4 mice). **(c**-**e)** Representative images of DS. Nuclei labeled with DAPI in blue. Dashed circles outline *Grpr+* cells. **(c)** FISH for *Chat* in green, *Grpr* in magenta, and *Ppp1r1b* in cyan. **(d)** FISH for *Drd2* in green, *Grpr* in magenta, and *Drd1* in cyan. **(e)** FISH for *Drd2* in green, *Grpr* in magenta, and *Penk* in cyan. **(f**-**j)** Mean ± SEM percentage of *Grpr*+ cells in the DS of WT mice that are **(f)** *Chat+* (n=7 mice), **(g)** *Ppp1r1b*+ (n=7 mice), **(h)** *Drd2*+ (n=12 mice), **(i)** *Drd1*+ (n=8 mice), or **(j)** *Penk*+ (n=8 mice). Related to Figure 2.

**Extended Data Fig. 5:**
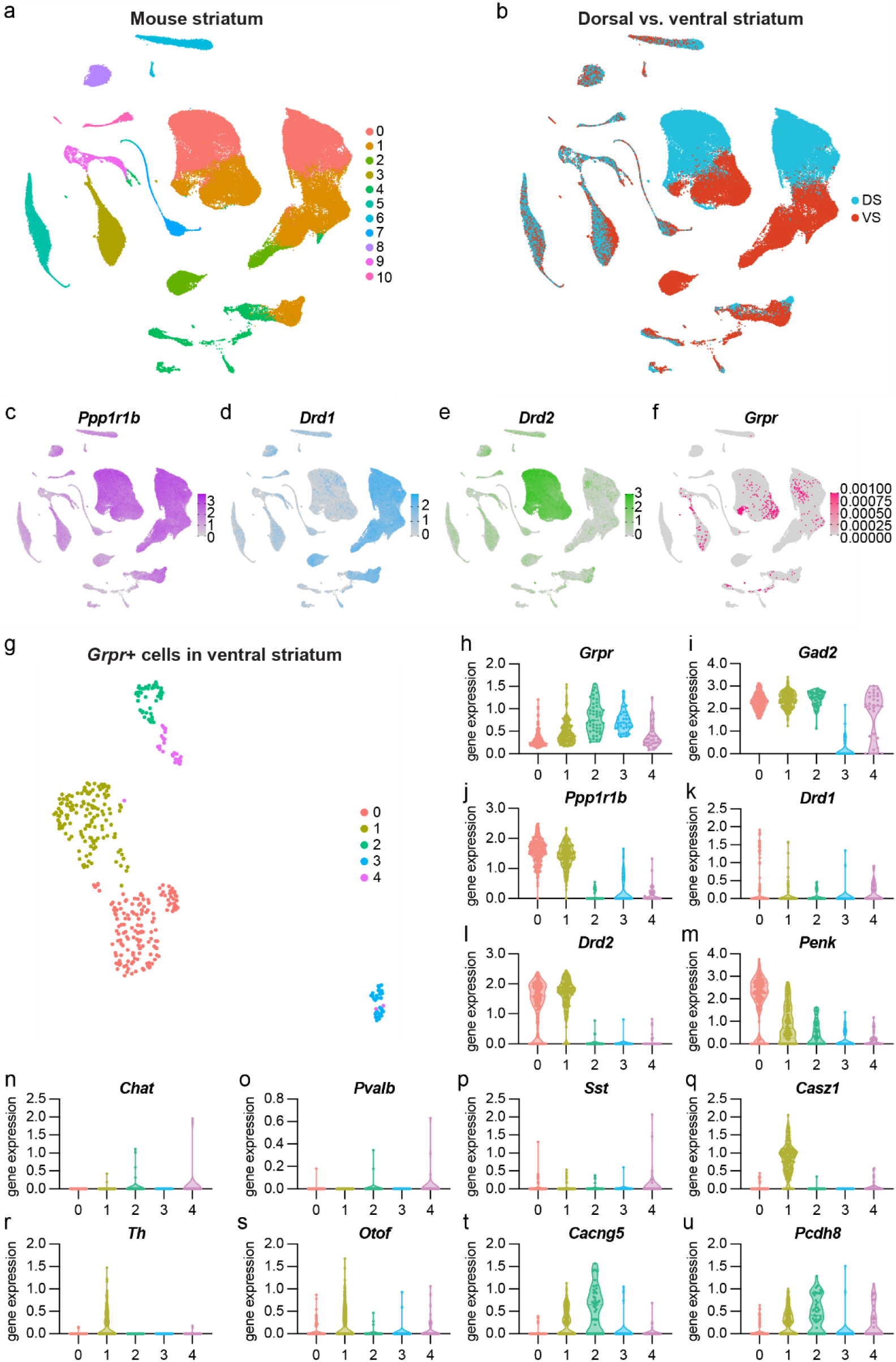
*Grpr* is expressed in multiple cell types in the mouse ventral striatum. **(a)** UMAP of mouse striatum single-cell RNAseq data, colored by cluster. Data and labels are from (Yao et al., 2023). Data were re-clustered at a resolution of 0.1. Clusters map broadly to cell types as follows: 0: CNU-LGE GABA, 1: CNU-LGE GABA, 2: astrocytes, 3: CNU-LGE GABA, 4: glutamatergic neurons, 5: OPC-Oligodendrocytes, 6: Immune, 7: OPC-Oligodendrocytes, 8: vascular, 9: DG-IMN Glut/OB-IMN GABA, 10: vascular. **(b)** UMAP of mouse striatum single-cell data from **a**, colored by dissection region: DS: dorsal striatum, VS: ventral striatum. **(c**-**f)** Feature plots of **(c)** *Ppp1r1b*, **(d)** *Drd1*, **(e)** *Drd2*, and **(f)** *Grpr* expression the mouse striatum. For **f** the color scale was adjusted to highlight any cell expressing *Grpr* transcripts. **(g)** UMAP of *Grpr*-expressing cells in the VS. All cells expressing any *Grpr* were subsetted, re-processed, and re-clustered at a resolution of 0.4. **(h**-**u)** Relative expression of **(h)** *Grpr*, **(i)** *Gad2*, **(j)** *Ppp1r1b*, **(k)** *Drd1*, **(l)** *Drd2*, **(m)** *Penk*, **(n)** *Chat*, **(o)** *Pvalb*, **(p)** *Sst*, **(q)** *Casz1*, **(r)** *Th***, (s)** *Otof*, **(t)** *Cacng5*, and **(u)** *Pcdh8* across the clusters identified in the UMAP of *Grpr-*expressing cells in **g**. For violin plots in **h**-**u**, each dot represents the expression level in a single cell. Related to Figure 2.

**Extended Data Fig. 6:**
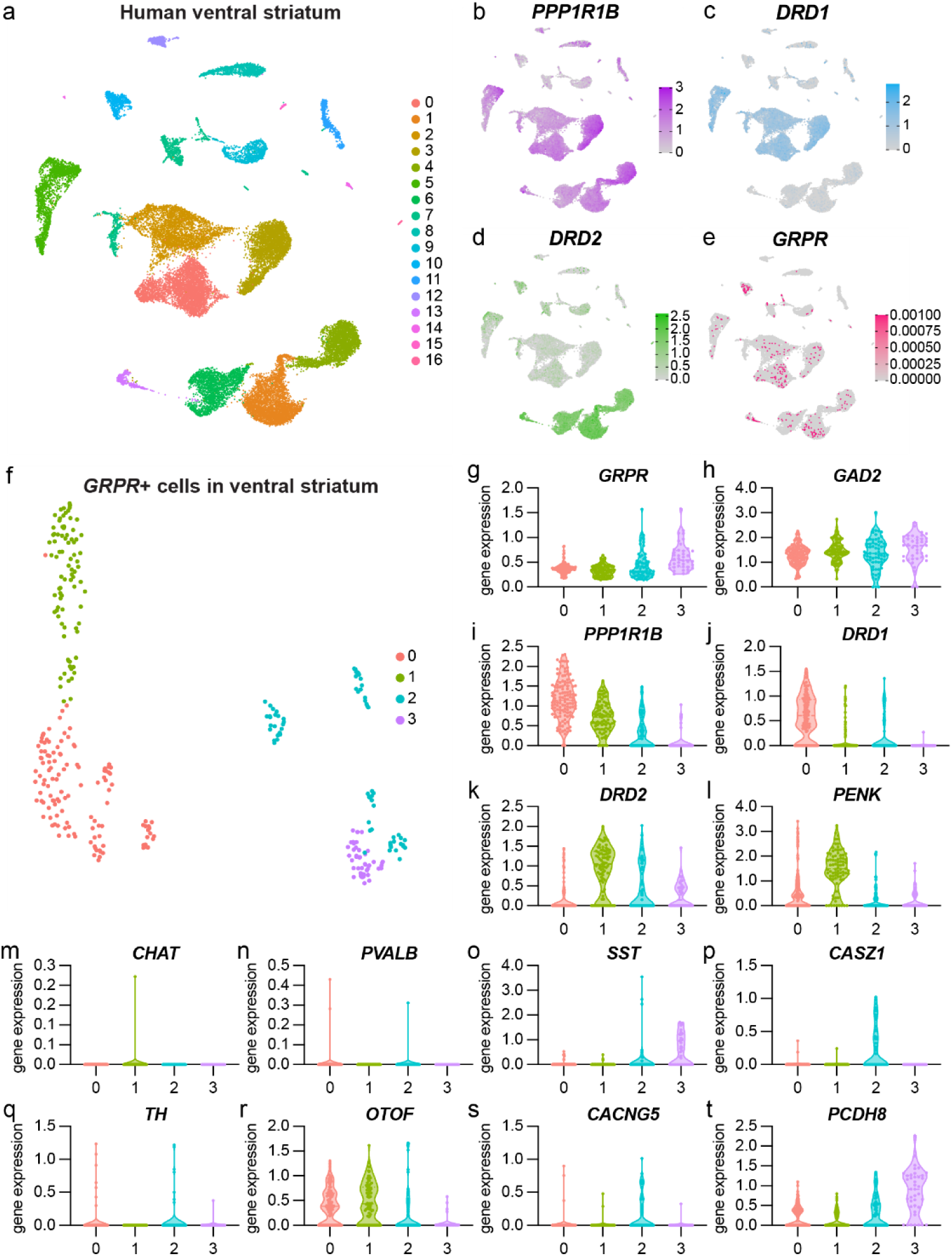
*GRPR* is expressed in multiple cell types in the human ventral striatum. **(a)** UMAP of human ventral striatum single-cell RNAseq data, colored by cell type annotation. Data and labels are from (Siletti et al., 2023). Clusters map broadly to cell types as follows: (0,1,2,3,4,6,13): Striatal projection neurons, 5: Eccentric striatal projection neurons, (7,9,10,16): splatter, 8: oligodendrocytes, 11: astrocytes, 12: oligodendrocyte Precursor, 14: microglia, 15: ependymal. **(b**-**e)** Feature plots of **(b)** *PPP1R1B*, **(c)** *DRD1*, **(d)** *DRD2*, and **(e)** *GRPR* expression the human striatum. For **e** the color scale was adjusted to highlight any cell expressing *GRPR* transcripts. **(f)** UMAP of *GRPR*-expressing cells in the ventral striatum. All cells expressing any *GRPR* were subsetted, re-processed, and re-clustered at a resolution of 0.3. **(g**-**t)** Relative expression of **(g)** *GRPR*, **(h)** *GAD2*, **(i)** *PPP1R1B*, **(j)** *DRD1*, **(k)** *DRD2*, **(l)** *PENK*, **(m)** *CHAT*, **(n)** *PVALB*, **(o)** *SST*, **(p)** *CASZ1*, **(q)** *TH***, (r)** *OTOF*, **(s)** *CACNG5*, and **(t)** *PCDH8* across the clusters identified in the UMAP of *GRPR-*expressing cells in **f**. For violin plots in **g**-**t**, each dot represents the expression level in a single cell. Related to Figure 2.

**Extended Data Fig. 7:**
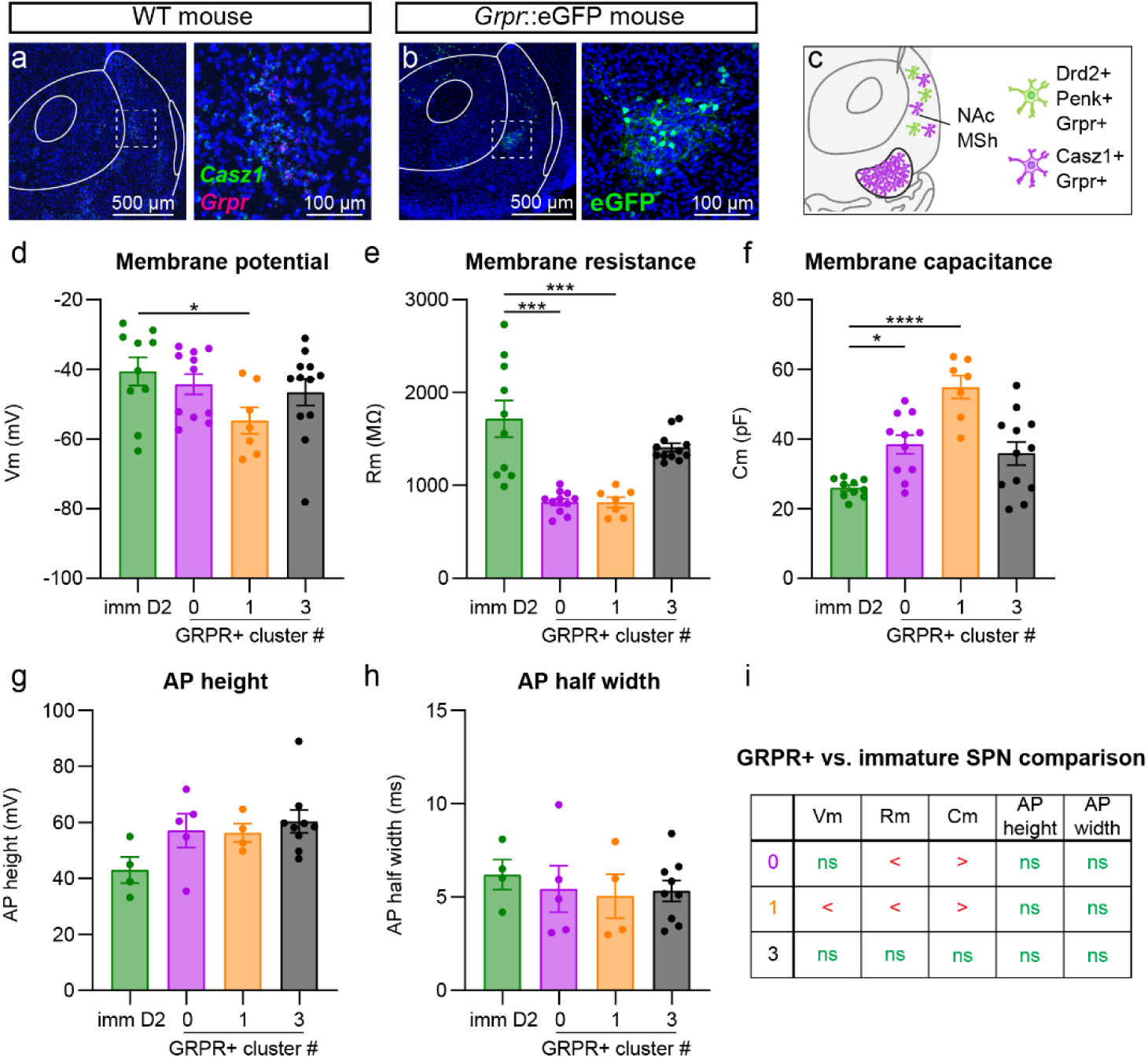
A subset of *Grpr+* cells have similar properties to immature SPNs in the NAc MSh. **(a)** Representative image of the NAc MSh. FISH for *Casz1* in green and *Grpr* in magenta. Nuclei labeled with DAPI in blue. Right panel is a zoom-in from the dashed box on the left. **(b)** Representative image of the NAc MSh from a *Grpr*::eGFP mouse. eGFP in green, DAPI labeled nuclei in blue. Right panel is a zoom-in from the dashed box on the left. **(c)** Summary schematic of NAc MSh showing three different populations of *Grpr*+ cells based on their genetic markers and location. **(d)** Mean ± SEM membrane potential of NAc MSh neurons. P8-P12 D2-GFP+ (imm D2) in dark green: n=10 cells from 6 mice, *Grpr*-eGFP+ cells in cluster 0 in purple: n=11 cells from 7 mice, *Grpr*-eGFP+ cells in cluster 1 in orange: n=7 cells from 5 mice, and *Grpr*-eGFP+ cells in cluster 3 in black: n=12 cells from n=8 mice (p=0.0837, one-way ANOVA; Kruskal-Wallis with Dunn’s correction for multiple comparisons, p>0.9999 imm D2 vs 0, *p=0.0336 imm D2 vs 1, and p=0.7357 4956 imm D2 vs 3). **(e)** Mean ± SEM membrane resistance of NAc MSh neurons. n is the same as for panel **d**. (****p<0.0001, one-way ANOVA; Kruskal-Wallis with Dunn’s correction for multiple comparisons, ***p=0.0002 imm D2 vs 0, ***p=0.0009 imm D2 vs 1, and p>0.9999 imm D2 vs 3). **(f)** Mean ± SEM membrane capacitance of NAc MSh neurons. n is the same as for panel **d**. (***p=0.0003, one-way ANOVA; Kruskal-Wallis with Dunn’s correction for multiple comparisons, *p=0.0380 imm D2 vs 0, ****p<0.0001 imm D2 vs 1, and p=0.1924 imm D2 vs 3). **(g)** Mean ± SEM AP height of NAc MSh neurons. P8-P12 D2-GFP+ (imm D2) in dark green: n=4 cells from 3 mice, *Grpr*-eGFP+ cells in cluster 0 (0) in purple: n=5 cells from 4 mice, *Grpr*-eGFP+ cells in cluster 1 (1) in orange: n=4 cells from 3 mice, and *Grpr*-eGFP+ cells in cluster 3 (3) in black: n=9 cells from n=5 mice. (p=0.1127, one-way ANOVA; Kruskal-Wallis with Dunn’s correction for multiple comparisons, p=0.1231 imm D2 vs 0, p=0.3821 imm D2 vs 1, and p=0.0609 imm D2 vs 3). **(h)** Mean ± SEM AP half-width of NAc MSh neurons. n is the same as for panel **g**. (p=0.6325, one-way ANOVA; Kruskal-Wallis with Dunn’s correction for multiple comparisons, p=0.7531 imm D2 vs 0, p=0.7586 imm D2 vs 1, and p>0.9999 imm D2 vs 3). **(i)** Summary table of **d**-**h**. Data for GRPR+ clustered cells in **d**-**h** are duplicated from Fig. 3 for comparison. Related to Figure 3.

**Extended Data Fig. 8:**
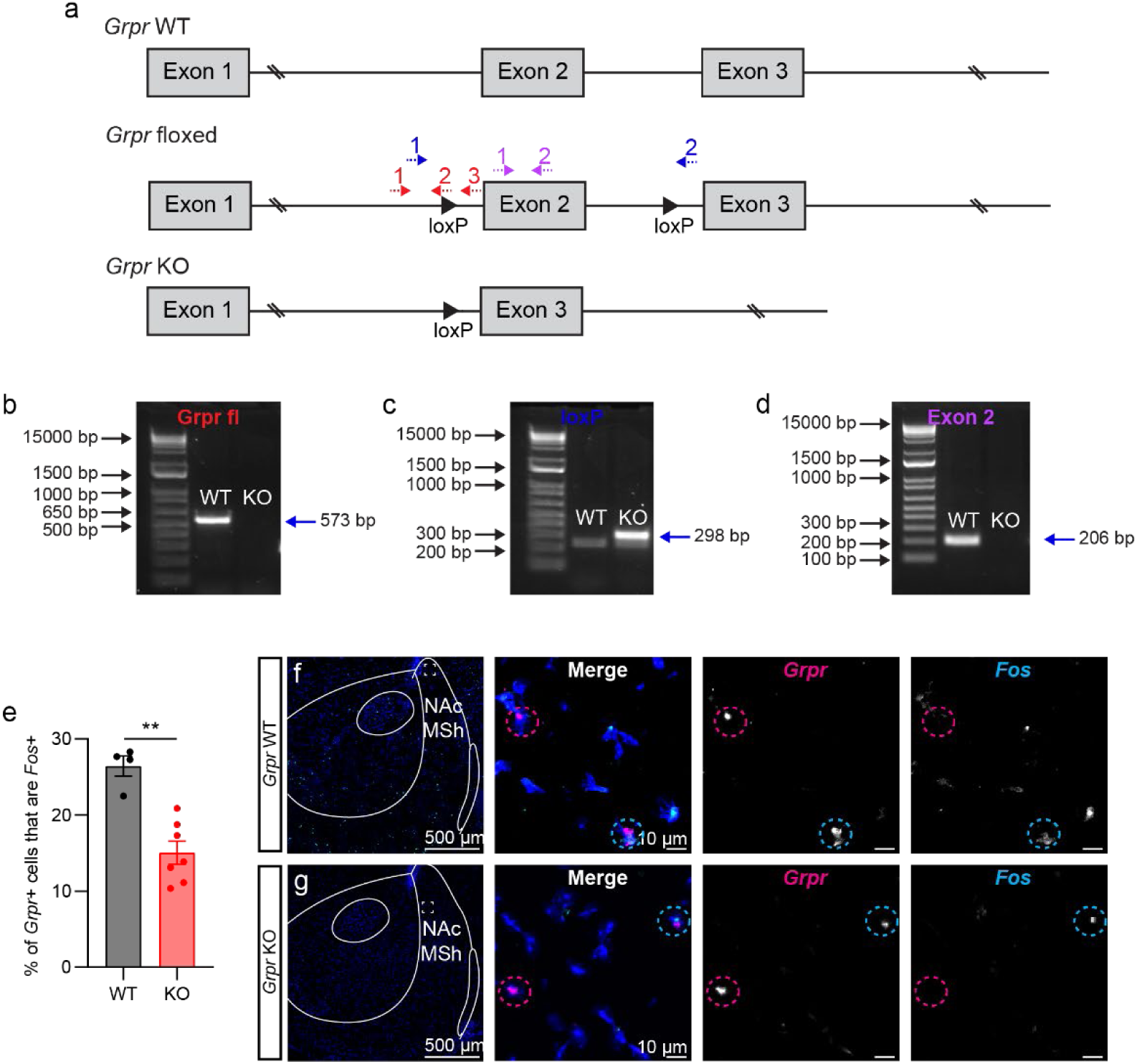
Mice lacking Grpr show reduced Fos expression in the NAc MSh. *Grpr* floxed mice were bred with CMV-Cre mice to delete *Grpr*. *Grpr* is on the X chromosome and only males were used for this analysis (*Grpr^fl/y^*). **(a)** Genotyping strategy for detection of the *Grpr* knock-out (KO), WT, and floxed alleles. Three primers (red) were designed to detect the presence of the loxP site within the floxed allele (1-WT forward, 2-floxed reverse, and 3-WT reverse). Two primers (blue) were designed to detect the loxP scar in the KO allele (1-loxP forward and 2-loxP reverse). Two primers (purple) were designed to detect exon 2 (1-Exon 2 forward and 2-Exon 2 reverse). **(b)** PCR detecting the presence of the floxed or WT allele for *Grpr*. WT mice were expected to have a band at 573 bp and KO mice to have no band. **(c)** PCR detecting the loxP scar, indicating successful recombination. WT mice were not expected to have a band as the sequence is too large to be amplified, while KO mice should have a band at 298 bp. **(d)** PCR for the presence of *Grpr* exon 2. WT mice were expected to have a band at 206 bp, while KO mice should not have a band. **(e)** Mean ± SEM percent *Grpr*+ cells in the NAc MSh that are *Fos*+ in *Grpr* WT (black) and *Grpr* KO (red) mice. (n=4 WT and n=7 KO mice, **p=0.0061, Mann-Whitney test). **(f**-**g)** Representative images of NAc MSh from **(f)** *Grpr* WT and **(g)** *Grpr* KO mice. FISH for *Grpr* in magenta and *Fos* in cyan. Dashed circles outline *Grpr*+ cells, magenta=*Grpr*+/*Fos*- and cyan=*Grpr*+/*Fos*+. Related to Figure 4.

**Extended Data Fig. 9:**
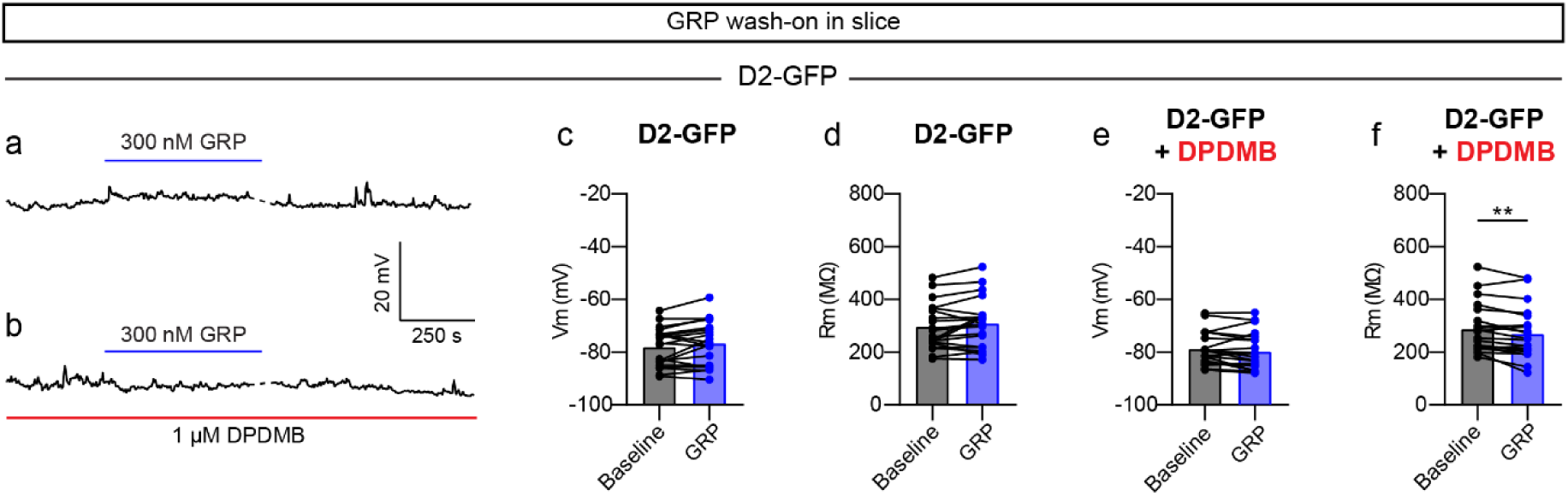
GRP does not increase the excitability of D2-expressing neurons in the NAc MSh. **(a)** Representative trace showing the membrane potential over time of a NAc MSh D2-GFP+ cell. Blue bar represents ten-minute wash-on of 300 nM GRP. **(b)** Representative trace showing the membrane potential over time of a NAc MSh D2-GFP+ cell. 300 nM GRP (blue bar) was washed-on in the presence of 1 μM DPDMB (red bar, entirety of recording). **(c)** Membrane potential of D2-GFP+ NAc MSh neurons at baseline (black) and after 300 nM GRP wash-on (blue); (n=20 cells from 20 mice; p=0.1054, Wilcoxon test). **(d)** Membrane resistance of D2-GFP+ NAc MSh neurons at baseline and after GRP wash-on; (n=20 cells from 20 mice; p=0.1429, Wilcoxon test). **(e)** Membrane potential of D2-GFP+ NAc MSh neurons at baseline and after GRP wash-on in the presence of DPDMB; (n=22 cells from 17 mice; p=0.0501, Wilcoxon test). **(f)** Membrane resistance of D2-GFP+ NAc MSh neurons at baseline and after GRP wash-on in the presence of DPDMB; (n=22 cells from 17 mice; **p=0.0025, Wilcoxon test). Related to Figure 6.

**Extended Data Fig. 10:**
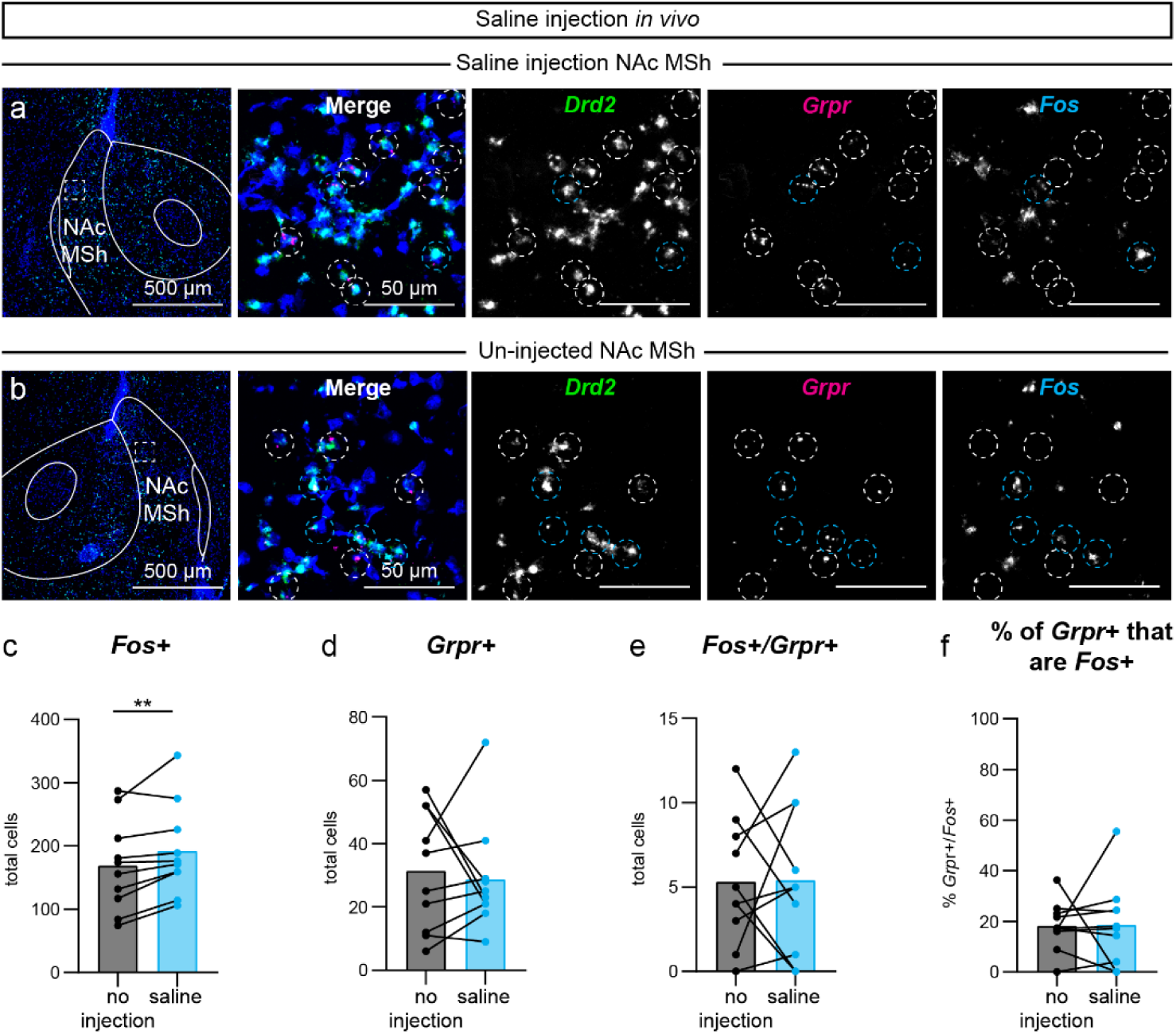
Saline injection into the NAc MSh does not induce *Fos* in *Grpr*- expressing cells. **(a-b)** Representative images of NAc MSh from the **(a)** saline-injected hemisphere and **(b)** un-injected hemisphere. FISH for *Drd2* in green, *Grpr* in magenta, and *Fos* in cyan. Nuclei are labeled with DAPI in blue. Dashed circles outline *Grpr*+ cells, white=*Grpr*+/*Fos*-, cyan=*Grpr*+/*Fos*+. **(c)** Total number of cells in the NAc MSh that were *Fos+* in the un-injected (black) vs saline-injected hemisphere (light blue) (**p=0.0098, Wilcoxon test). **(d)** Total number of cells in the NAc MSh that were *Grpr+* in the un-injected vs saline-injected hemisphere (p=0.9414, Wilcoxon test). **(e)** Total number of *Grpr*+ cells in the NAc MSh that were *Fos+* in the un-injected vs saline-injected hemisphere (p=0.9570, Wilcoxon test). **(f)** Percentage of *Grp*r+ cells in the NAc MSh that were *Fos*+ in the un-injected vs saline-injected hemisphere (p=0.7695, Wilcoxon test). For panels **c**-**f** data are summed from two coronal sections per mouse (n=10 mice). Related to Figure 6.

**Extended Data Fig. 11:**
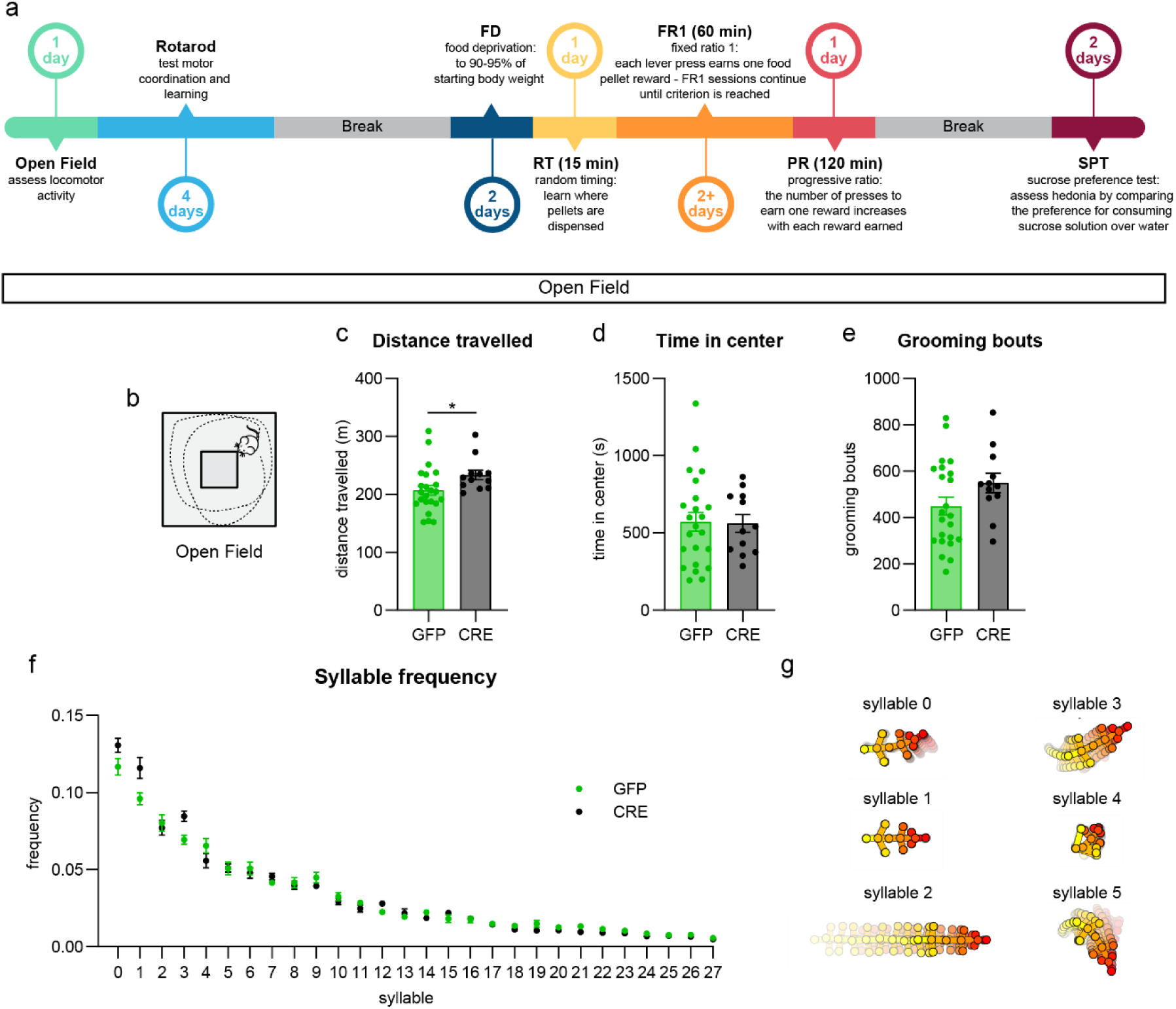
Mice lacking *Grpr* in the NAc MSh have increased locomotor activity but no differences in behavioral syllable usage the open field. **(a)** Timeline of behavioral assays in *Grpr* cKO and littermate control mice. **(b)** Schematic of the open field. **(c)** Mean ± SEM total distance travelled in the one-hour session for control (GFP injected, green) and *Grpr* cKO (CRE injected, black). Mann-Whitney test, *p=0.0234 (n=23 GFP and 12 CRE mice for all panels). **(d)** Mean ± SEM total time spent in the center of the open field during the one-hour session. Center was defined as a 20 cm x 20 cm square in the middle of the arena. Mann-Whiney test, p=0.8506. **(e)** Mean ± SEM total number of grooming bouts during the one-hour open field session. Mann-Whitney test, p=0.1723. **(f)** Mean ± SEM frequency of syllable usage for control (GFP injected, green) and *Grpr* cKO (CRE injected, black) mice. Syllable descriptions: 0-left turn, 1-static, 2-forward, 3-left turn 2, 4-grooming, 5-right turn, 6-forward 2, 7-left turn 3, 8-forward left, 9-right turn 2, 10-forward 3, 11-grooming 2, 12-grooming 3, 13-grooming 4, 14-backward right, 15-backward left, 16-forward 4, 17-left turn 4, 18-grooming 5, 19-forward 5, 20-forward 6, 21-forward 7, 22-backward, 23-forward 8, 24-forward right, 25-right turn 3, 26-grooming 6, 27-left turn 5. Kruskal-Wallis with Dunn’s correction for multiple comparisons, 0: p=0.1724,1: p=0.1540, 2: p=0.5629, 3: p=0.1498, 4: p=0.2520, 5: p=0.8470, 6: p=0.6170, 7: p=0.2589, 8: p=0.7552, 9: p=0.4081, 10: p=0.5152, 11: p=0.2021, 12: p=0.1498, 13: p=0.8276, 14: p=0.1540, 15: p=0.2021, 16: p=0.7552, 17: p=0.6745, 18: p=0.1782, 19: p=0.1540, 20: p=0.5186, 21: p=0.2021, 22: p=0.2180, 23: p=0.5214, 24: p=0.1540, 25: p=0.5629, 26: p=0.4716, 27: p=0.2180. **(g)** Average pose trajectories for the top six Keypoint-MoSeq syllables. Each trajectory includes ten poses, starting 165 ms before and ending 500 ms after syllable onset. Related to Figure 7.

**Extended Data Fig. 12:**
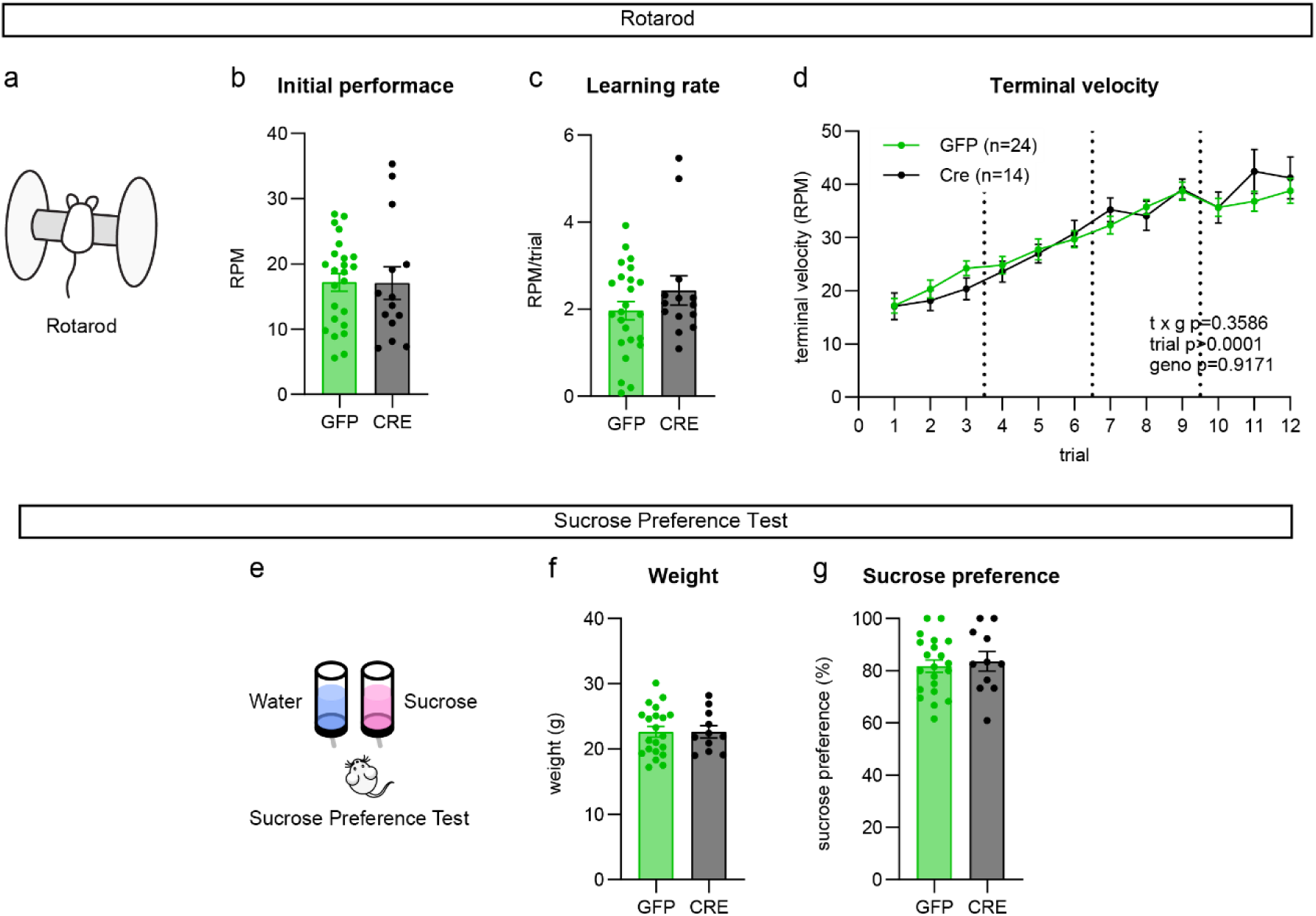
Mice lacking *Grpr* in the NAc MSh show no differences in the rotarod or sucrose preference tests. **(a)** Schematic of the accelerating rotarod. **(b)** Mean ± SEM performance on trial 1 of the rotarod in control (GFP-injected, green) vs *Grpr* cKO (CRE-injected, black) mice. Mann-Whitney test, p=0.6700 (n=24 GFP and 14 CRE mice). **(c)** Mean ± SEM of learning rate, measured as the slope of the linear regression of the performance per mouse across all trials. Mann-Whitney test, p=0.5201. (n is the same as in panel **b**) **(d)** Mean ± SEM rotarod performance for GFP-injected and CRE-injected mice across all trials, measured as terminal velocity. Two-way repeated measures ANOVA p-values are shown in the graph. (n is the same as for panel **b**). **(e)** Schematic of sucrose preference test. **(f)** Mean ± SEM weight of GFP-injected and CRE-injected mice just prior to the sucrose preference test. Mann-Whitney test, p=0.9300 (n=21 GFP and 11 CRE mice). **(g)** Mean ± SEM sucrose preference measured as the percentage of total liquid consumed that was 5% sucrose. Welch’s two-tailed t-test, p=0.6785; (n is the same as for panel **f**). Related to Figure 7.

**Extended Data Fig. 13:**
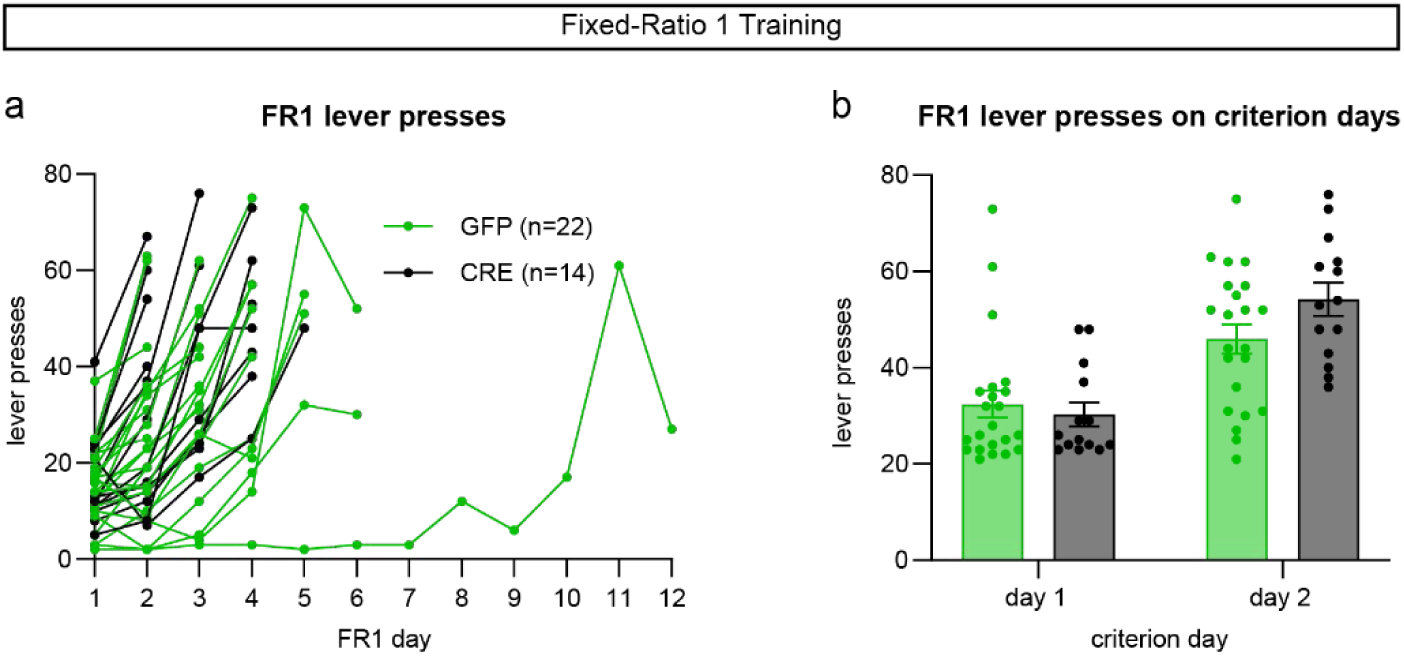
Mice lacking *Grpr* in the NAc MSh have normal performance in fixed ratio training sessions. **(a)** Number of lever presses on each day of the fixed-ratio 1 (FR1) training for individual mice. Control (GFP-injected) mice are shown in green and *Grpr* cKO (CRE-injected) mice are shown in black. (n=22 GFP and 14 CRE mice for all panels) **(b)** Mean ± SEM total lever presses on the final two days of FR1 training in control (GFP-injected, green) and *Grpr* cKO (CRE-injected, black) mice. Two-way repeated measures ANOVA: day x genotype p=0.0671, day ****p<0.0001, genotype p=0.3845. Holm-Sidak’s multiple comparisons test, day 1: p=0.8596, day 2: p=0.1278. Related to Figure 7.

